# A cell atlas of human adrenal cortex development and disease

**DOI:** 10.1101/2022.12.13.520231

**Authors:** Ignacio del Valle, Matthew D Young, Gerda Kildisiute, Olumide K Ogunbiyi, Federica Buonocore, Ian C Simcock, Eleonora Khabirova, Berta Crespo, Nadjeda Moreno, Tony Brooks, Paola Niola, Katherine Swarbrick, Jenifer P Suntharalingham, Sinead M McGlacken-Byrne, Owen J Arthurs, Sam Behjati, John C Achermann

**Affiliations:** Genetics and Genomic Medicine Research and Teaching Department, UCL Great Ormond Street Institute of Child Health, University College London, London, WC1N 1EH, UK; Wellcome Sanger Institute, Wellcome Genome Campus, Hinxton, Cambridge, CB10 1SA, UK; Department of Histopathology, Great Ormond Street Hospital for Children NHS Foundation Trust, London, WC1N 3JH, UK; Developmental Biology and Cancer Research and Teaching Department, UCL Great Ormond Street Institute of Child Health, University College London, London, WC1N 1EH, UK; Department of Clinical Radiology, Great Ormond Street Hospital for Children NHS Foundation Trust, London, WC1N 3JH, UK; Population, Policy and Practice Research and Teaching Department, UCL Great Ormond Street Institute of Child Health, University College London, London, WC1N 1EH, UK; UCL Genomics, Zayed Centre for Research, UCL Great Ormond Street Institute of Child Health, University College London, London, WC1N 1DZ, United Kingdom; Cambridge University Hospitals NHS Foundation Trust, Cambridge, CB2 0QQ, UK; Department of Paediatrics, University of Cambridge, Cambridge, CB2 0QQ, UK

**Author notes:** Corresponding author Correspondence: John C. Achermann, MB MD PhD, Genetics & Genomic Medicine Research and Teaching Department, UCL Great Ormond Street Institute of Child Health, University College London, London, WC1N 1EH, UK. Jointly supervised the work (Sam Behjati, John C Achermann).

**Keywords:** Adrenal gland, transcriptomics, scRNA-seq, steroidogenesis, HOPX, RSPO3, imprinted genes, adrenal insufficiency

## Abstract

The adrenal glands synthesize and release essential steroid hormones such as cortisol and aldosterone, but the mechanisms underlying human adrenal gland development are not fully understood. Here, we combined single-cell and bulk RNA-sequencing, spatial transcriptomics, immunohistochemistry and micro-focus computed tomography to investigate key aspects of adrenal development in the first 20 weeks of gestation. We demonstrate rapid adrenal growth and vascularization, with cell division in the outer definitive zone (DZ). Steroidogenic pathways favor androgen synthesis in the central fetal zone (FZ), but DZ capacity to synthesize cortisol and aldosterone develops with time. Core transcriptional regulators were identified, with a role for HOPX in the DZ. Potential ligand-receptor interactions between mesenchyme and adrenal cortex were seen (e.g., *RSPO3*/*LGR4*). Growth-promoting imprinted genes were enriched in the developing cortex (e.g. *IGF2, PEG3*). These findings reveal new aspects of human adrenal development, and have clinical implications for understanding primary adrenal insufficiency and related postnatal adrenal disorders, such as adrenal tumor development, steroid disorders and neonatal stress.

## Introduction

The mature, adult adrenal glands are essential endocrine organs that consist of an outer cortex and a central medulla. The adrenal cortex has three layers that synthesize and release key groups of steroid hormones^1–4^. Mineralocorticoids (e.g., aldosterone) are released from the outer zona glomerulosa and are needed for salt retention and blood pressure maintenance. Glucocorticoids (e.g. cortisol) are released predominantly from the zona fasciculata and are needed for wellbeing and glucose regulation. Weak androgens (e.g., dehydroepiandrosterone) are released from the inner zona reticularis and influence adrenarche in mid-childhood, with potential effects on health in adult women^5–7^. In contrast, the central adrenal medulla is neuroectodermal in origin and releases epinephrine (adrenaline) and norepinephrine (noradrenaline)^8^. Thus, the adrenal glands play an essential role in the acute stress response, many aspects of physiological homeostasis, and long-term wellbeing.

Disruption of adrenal gland function (known as primary adrenal insufficiency, PAI) leads to glucocorticoid insufficiency, often combined with mineralocorticoid insufficiency^9–11^. PAI can present at various ages with symptoms such as malaise, weight loss, hyperpigmentation and hypotension, and can be fatal if not diagnosed and treated appropriately^9^. Although autoimmune destruction of the adrenal gland (sometimes referred to as “Addison disease”) is the most common cause of PAI in adolescents and adults, around 30 different single gene disorders have now been identified that result in PAI through diverse processes such as impaired development (hypoplasia), blocks in steroid biosynthesis (congenital adrenal hyperplasia, CAH), adrenocorticotropic hormone (ACTH) resistance (‘familial glucocorticoid deficiency’, FGD), and metabolic conditions^10, 12, 13^. PAI often presents soon after birth, or more gradually in childhood or even adulthood. Individuals with PAI require lifelong adrenal steroid hormone replacement, with management sometimes modified based on the underlying cause^9, 14^.

In humans, the adrenal cortex develops from bilateral condensations of intermediate mesoderm, known as the “adrenogonadal primordium”, at approximately 4 weeks post conception (wpc) (6 weeks gestation)^15–18^. These structures are in close proximity to the developing kidneys, and give rise to both the adrenal gland and gonad (testis, ovary)^15, 19^. The adrenal cortex and gonad share several distinct functional pathways, such as the ability to synthesize steroid hormones and regulation by the nuclear receptor, NR5A1 (also known as steroidogenic factor-1, SF-1)^20–23^. In contrast, the adrenal medulla is ectodermal in origin and arises from Schwann cell precursor cells that migrate into the adrenal gland and differentiate into sympathoblastic and chromaffin cells^8, 24^. These cells ultimately coalesce centrally to form the adrenal medulla.

Although insights into adrenal development and function are being obtained from studies in model systems (e.g., mice, zebrafish)^25–29^, the adrenal cortex in humans and higher primates has distinct structural and functional components^30^. Most notable is the development of a large fetal zone (FZ), which is capable of synthesizing and releasing substantial amounts of the weak androgen, dehydroepiandrosterone (DHEA) and its sulfated form, DHEA-S. DHEA is converted to estrogens by the placenta, which enter the maternal circulation during pregnancy^16^. The FZ regresses in the first few months of postnatal life^30, 31^. The teleological function of the FZ is not known although DHEA may have a role in neurodevelopment^32^.

Mice have an X-zone that regresses with sexual maturity (males) orpregnancy (females), but similarities with the human FZ are somewhat limited^33–35^. Furthermore, cortisol is the primary glucocorticoid synthesized by the adrenal gland in humans whereas rodents generate higher concentrations of corticosterone^3^.

In recent years, a limited number of studies of human adrenal development or fetal adrenal steroidogenesis have been undertaken using gene expression approaches or focussed RT- PCR/immunohistochemistry (IHC) of steroid pathways^18, 20, 36–40^. However, few data currently exist for detailed transcriptomic analysis of the human adrenal cortex at a single cell level during the critical first half of gestation (to 20wpc) or transcriptomics linked to developmental anatomy. We therefore developed a multimodal approach to investigate human adrenal cortex development in detail.

## Results

### The developing adrenal gland has a defined transcriptomic profile

In order to study the key biological events involved in human adrenal development, we integrated single-cell RNA-sequencing (scRNA-seq), bulk RNA-seq, spatial transcriptomics, immunohistochemistry (IHC) and micro-focus computed tomography (micro-CT) imaging across a critical developmental time-frame between 6 to 20wpc (Fig. 1a, Supplementary Data 1).

**Fig. 1.**
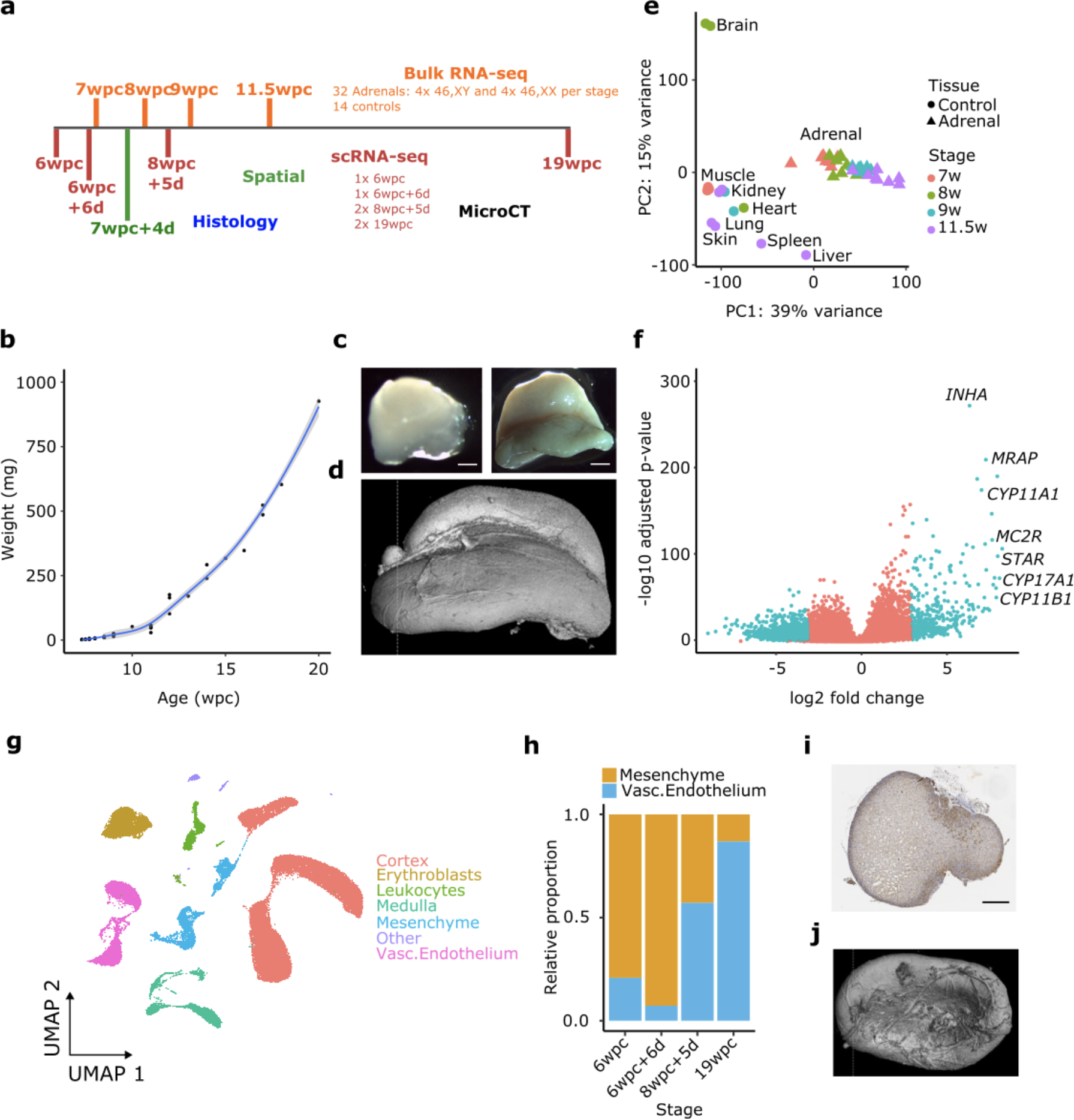
Study design, adrenal development and transcriptome analysis. **a** Overview of the study design for generating bulk transcriptomes (bulk RNA-seq), single-cell mRNA transcriptomes (scRNA-seq), spatial transcriptomics, microCT (micro-focus computed tomography) and histology/immunohistochemistry. Stages are shown as weeks (w) and days (d) post-conception (pc). **b** Growth curve of the adrenal gland between 7 weeks post-conception and 2 days (7wpc+2d) and 20wpc (*n* = 36). Data for single glands are shown. **c** Photographs of adrenal glands (10% formalin) at 6wpc+6d (left, scale bar 300μm) and 16 wpc (right, scale bar 3mm) to show marked growth and anatomical changes. **d** MicroCT surface image of the adrenal gland at 17wpc showing the anterior sulcus and vasculature. **e** Principal component analysis (PCA) of bulk transcriptome data for adrenal glands at 7wpc (*n* = 8), 8wpc (*n* = 8), 9wpc (*n* = 8) and 11.5wpc (*n* = 8), and control tissues (*n* = 14, from 8 different tissues) as indicated. **f** Volcano plot showing differential gene expression of genes in the bulk transcriptome adrenal gland dataset (total *n* = 32) compared to controls (*n* = 14). Selected highly differentially expressed adrenal genes are indicated. **g** UMAP of scRNA-seq transcriptome data from four adrenal glands (6w, 6wpc+6d, 8wpc+5d, 19w) with the major different cell populations annotated (6wpc, *n* = 3047 cells; 6wpc+6d, *n* = 2650 cells; 8wpc+5d, *n* = 23313 cells; 19wpc, *n* = 15348 cells). **h** Relative proportion of mesenchyme and vascular endothelial cells in the adrenal gland at each time point studied. **i** Section of the adrenal gland at 8.5wpc stained for vascular endothelial growth factor receptor 1 (VEGFR1) expression. Scale 400μm. **j** MicroCT (17wpc) to show the extensive surface vascular network on the inferior surface of the gland.

During this period, the adrenal gland undergoes rapid growth, and specific morphological changes such as the development of a deep sulcus and marked increases in vascularization (Fig. 1b-d, Supplementary Fig. 1, Supplementary Movie 1).

At a global level (bulk RNA-seq), the developing adrenal gland showed a well-defined transcriptomic profile compared to control tissues. This transcriptome is more similar to kidney at early stages of development (7wpc) but becomes increasingly distinct with age (Fig. 1e, Supplementary Fig. 2). A subset of highly differentially expressed adrenal-specific genes was identified, including known genes (e.g. *MC2R*, *STAR*, *CYP11A1*) as well as several genes not previously identified as differentially-expressed in adrenal development (e.g.*CLRN1*, *MIR202HG, FAM166B*) (Fig. 1f, Supplementary Fig. 2, Supplementary Data 2).

In order to define specific cell populations within the adrenal gland in more detail, single cell mRNA transcriptome analysis (scRNA-seq) was undertaken at four timepoints (6wpc, 6wpc+6days (d), 8wpc+5d, 19wpc) (Fig. 1a, g)^24^. This analysis clearly identified a cluster of adrenal cortex cells, with strong enrichment for genes involved in steroidogenesis (Fig. 1g, Fig. 2a, Supplementary Data 3). Other major clusters included cells that contribute to the developing adrenal medulla (Schwann cell precursors, sympathoblastic cells, chromaffin cells and recently described “medullary bridge” cells)^24^, as well as mesenchymal cells, vascular endothelial cells, erythroblasts and leukocytes (Fig. 1g, Supplementary Fig. 3). The relative proportion of mesenchymal cells decreased over time with differentiation, whereas the vascular endothelial components and erythroblast cells increased, consistent with the development of an extensive vascular network supplying the adrenal gland and a network of sinusoids within it, necessary for the release of steroid hormones into the developing fetal circulation (Fig. 1h-j, Supplementary Fig. 3).

**Fig. 2.**
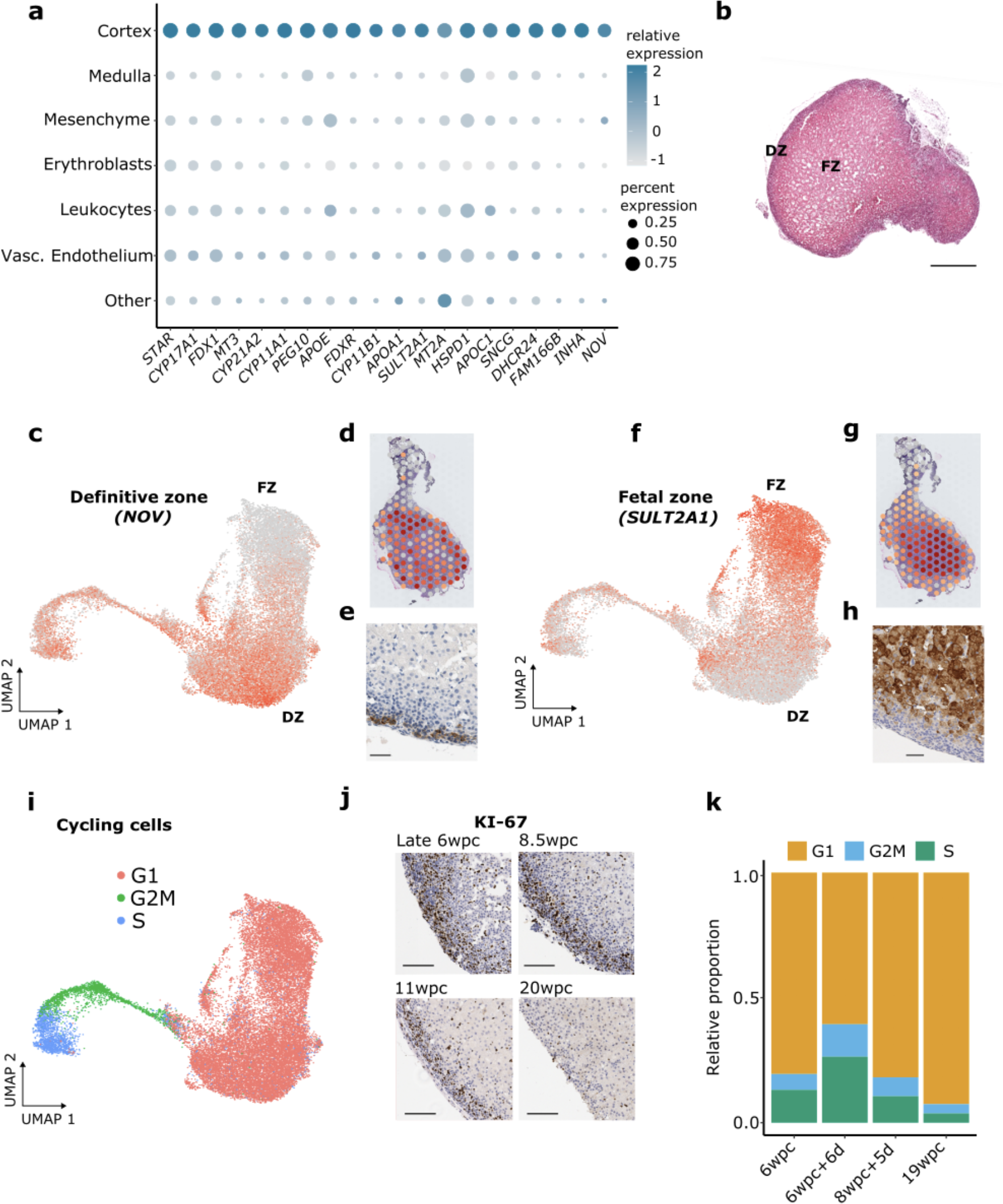
Adrenal cortex zonation and proliferation. **a** Dot plot to show the most highly differentially-expressed genes in the adrenal cortex single cell transcriptome (scRNA-seq) compared to other cells in the adrenal gland. **b** Histology of the human fetal adrenal gland at 8.5wpc (H&E staining). DZ, definitive zone; FZ, fetal zone. Scale 400μm. **c-e** The developing definitive zone shown by *NOV* (also known as *CCN3*) expression using a single cell mRNA transcriptome UMAP (**c**), spatial transcriptomic spotplot (7wpc+4d, darker red shows higher expression) (**d**) and immunohistochemistry (11wpc; scale 50μm) (**e**). Integrated data from samples at all four time points are shown. **f-h** The developing fetal zone shown by *SULT2A1* expression using a single cell mRNA transcriptome UMAP (**f**), spatial transcriptomic spotplot (7wpc+4d) (**g**) and immunohistochemistry (11wpc; scale 50μm) (**h**). **i** Integrated UMAP showing cell cycle states. **j** Immunohistochemistry of fetal adrenal gland showing KI67 expression as a marker of cell proliferation at different ages. Scales all 100μm. **k** Relative proportion of cells in each cell cycle state at each time point.

### The adrenal cortex has distinct zones

Subsequent analysis focussed on the fetal adrenal cortex (Fig. 1g, Fig. 2a, Supplementary Fig. 4), as relatively few data are available for cortex development in humans, especially in the second trimester, and single cell mRNA transcriptome analysis allows new insights to be obtained.

Histologically, the fetal adrenal cortex is broadly divided into an outer definitive zone (DZ), somewhat similar to the postnatal zona glomerulosa and zona fasciculata, and an inner fetal zone (FZ) consisting of large cytomegalic cells interspersed with vascular sinusoids (Fig. 2b). A distinct capsule forms around the adrenal gland during the first trimester, with a putative transition zone developing later in the second trimester^30^.

In order to study cortex zonation in more detail, we used nephroblastoma overexpressed/cysteine-rich protein 61/connective tissue growth factor/nephroblastoma overexpressed gene-3 (*NOV*, also known as *CCN3*) as a marker for the DZ, and sulfotransferase 2A1 (*SULT2A1*) as a marker for the FZ^39, 41, 42^. These genes differentiated the DZ and FZ clearly in an integrated scRNA-seq dataset, as well as by spatial transcriptomics (7wpc+4d) and IHC (11 wpc data shown) (Fig. 2c-h). Of note, cycling cells (S phase, GM2 phase) clustered more closely with the DZ rather than the FZ in all stages studied (Fig. 2i, Supplementary Fig. 5a-c). This finding was supported by IHC using KI-67 as a marker of cell division (Fig. 2j). The relative proportion of dividing cells was highest during early development (Fig. 2j, k), consistent with rapid growth of the gland during this time (Fig. 1c, Supplementary Fig. 1). During the earliest stage (6w), a trajectory of cells from the DZ to FZ was seen (Supplementary Fig. 5d). Taken together, these data suggest that the DZ is a more active region of cell division compared to the FZ, and with potential centripetal cell differentiation at least in early development^43–46^.

### Fetal adrenal steroidogenesis favors DHEA synthesis

A major role of the mature, postnatal adrenal cortex is to synthesize and release steroid hormones such as cortisol and aldosterone. The extent to which the developing adrenal gland has the biosynthetic capacity to produce these steroids is still unclear. It is well established that the FZ synthesizes and releases large amounts of DHEA(S), due to a relative lack of the enzyme 3 β-hydroxysteroid dehydrogenase type 2 (3β-HSD2, encoded by *HSD3B2*) and likely high expression of genes encoding enzymes needed for androgen biosynthesis (i.e. *CYP17A1*, *POR*, *CYB5A*). Although a transient wave of *HSD3B2*/3β-HSD2 expression has been shown at around 8-9 wpc^37, 39^, evidence is still emerging as to when the necessary enzymes for glucocorticoid (e.g., cortisol) and mineralocorticoid (e.g., aldosterone) synthesis are expressed during human adrenal development, especially into the second trimester^38, 39^.

In order to explore this further, we analyzed time-series bulk RNA-seq data (between 7wpc and 11.5wpc), which showed a clear temporal increase in expression of the ACTH receptor (*MC2R*), as well as steroidogenic acute regulator protein (STAR) and most other steroidogenic enzymes (Fig. 3a, b, Supplementary Fig. 6). These data show that the machinery for ACTH-dependent cholesterol processing is in place during early development, and increases with age.

**Fig. 3.**
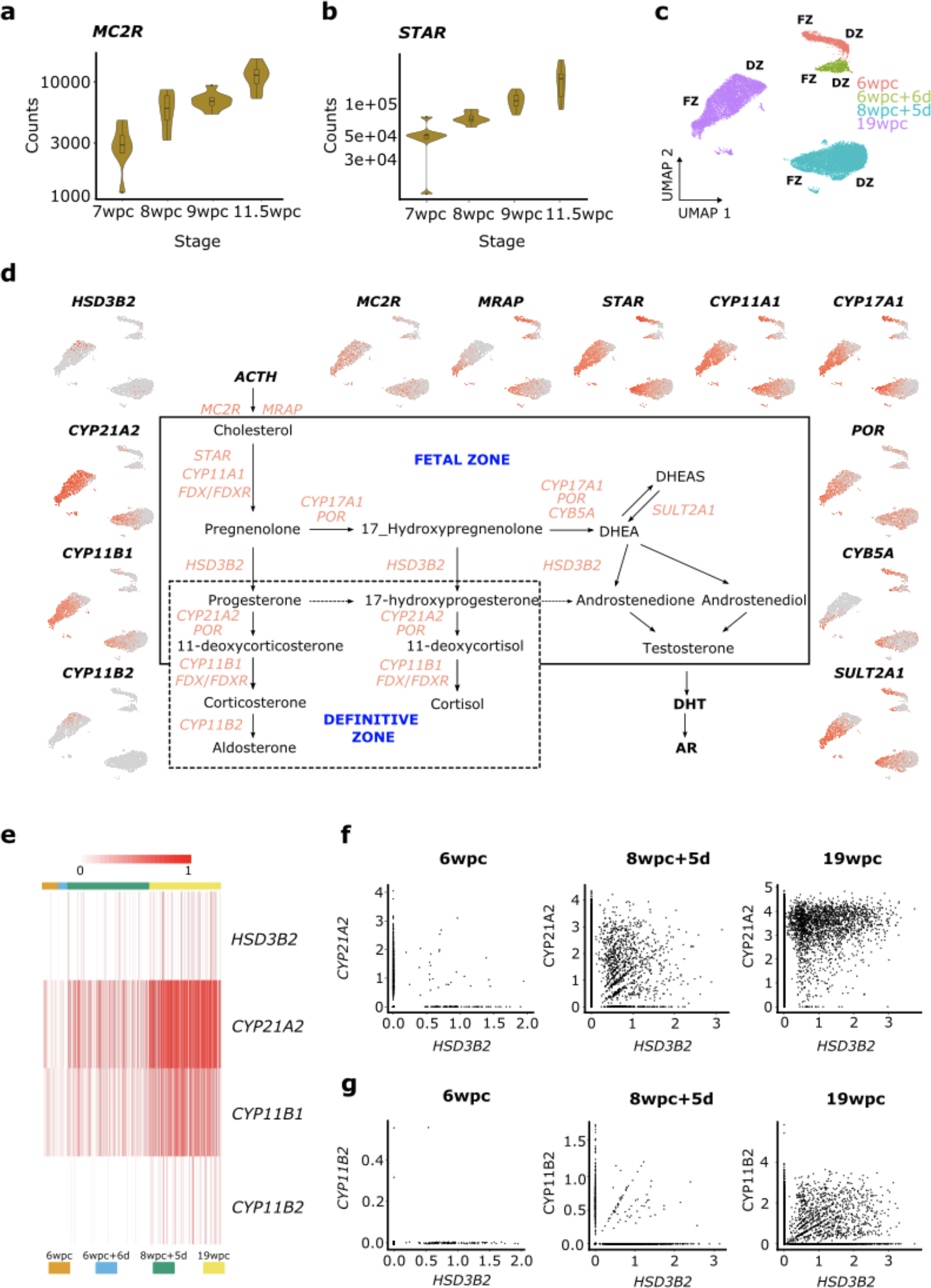
Expression of classic steroidogenic pathway genes during human adrenal development. **a** Time-series bulk RNA-seq expression (normalized counts) of the melanocortin-2 receptor gene (*MC2R*), encoding the adrenocorticotropin (ACTH) receptor. **b** Time-series bulk RNA-seq expression of the gene encoding steroidogenic acute regulatory protein (*STAR*). **c** UMAP of adrenal cortex clusters used for subsequent analysis. DZ, definitive zone; FZ, fetal zone. **d** Graphical representation of the “classic” steroidogenic pathway showing the key genetic components leading to the synthesis of mineralocorticoids (e.g. aldosterone), glucocorticoids (e.g. cortisol) and androgens (e.g. dehydroepiandrosterone (DHEA), androstenedione, testosterone). Feature plots showing the expression of key genes in the adrenal cortex clusters at different time points are shown. ACTH, adrenocorticotropin; AR, androgen receptor; *CYB5A*, cytochrome 5A; *CYP11A1*, P450 side-chain cleavage enzyme; *CYP11B1*, 11β-hydroxylase; *CYP11B2*, aldosterone synthase; *CYP17A1*, 17⍺-hydroxylase/17,20-lyase; *CYP21A2*, 21-hydroxylase; DHEA(S), dehydroepiandrosterone (sulfate); DHT, dihydrotestosterone; *HSD3B2*, 3β-hydroxysteroid dehydrogenase type 2; *MC2R*, melanocortin-2 receptor (ACTHR); *MRAP*, MC2R-accessory protein; *POR*, P450 oxidoreductase; *STAR*, steroidogenic acute regulatory protein; *SULT2A1*, sulfotransferase 2 A1. **e** Heatmap of scRNA-seq expression of *HSD3B2*, *CYP21A2*, *CYP11B1* and *CYP11B2* at different ages in the adrenal cortex clusters. **f** Scatter plots of expression of *HSD3B2* in individual cortex single cells (scRNA-seq) compared to *CYP21A2* at three different time points (6wpc, 8wpc+5d, 19wpc). **g** Scatter plots of expression of *HSD3B2* in individual cortex single cells (scRNA-seq) compared to *CYP11B2* at three different time points (6wpc, 8wpc+5d, 19wpc).

Next, a scRNA-seq dataset was generated sub-setting the annotated adrenal cortex cells at each time point studied, with cycling cells removed (see Uniform Manifold Approximation and Projection (UMAP), Fig. 3c). Across all stages, cells of the FZ region showed high expression of genes encoding the key enzymes needed for DHEA synthesis (*STAR*, *CYP11A1*, *CYP17A1*, *POR*, *CYB5A*) as well as of *SULT2A1* (encoding sulfotransferase 2A1), which is required for sulfation of DHEA to DHEA-S and protects the developing fetus from androgen exposure (Fig. 3d). As expected, *HSD3B2* expression was low in the FZ during development, resulting in the likely shuttling of steroid precursors (e.g. pregnenolone) into the androgen pathway. The high expression of *STAR* and *CYP11A1* in the FZ cluster was mirrored by high expression of the ACTH receptor and its accessory protein (*MRAP*) suggesting not only that the FZ has the capacity to be highly biosynthetically active but also that FZ DHEA synthesis may be ACTH-dependent. Of note, enzymes proposed to be involved in the “backdoor” pathway of androgen synthesis^40^ were not strongly expressed, although several components of the pathway needed for 11-oxygenation of androgens^47^ were (Supplementary Figs. 7-9).

As 3β-hydroxysteroid dehydrogenase type 2/*HSD3B2* is effectively a “gatekeeper” to glucocorticoid and mineralocorticoid biosynthesis (Fig. 3d), we investigated *HSD3B2* expression across time-series data. Although a potential transiently higher expression was seen at 8wpc in bulk RNA-seq data (Supplementary Fig. 6), as reported previously^37, 39^, single-cell transcriptomic data showed overall greater increase in *HSD3B2* across time, with the highest levels in the DZ cluster at 19wpc (Fig. 3e). A similar graded increase in *CYP21A2* (encoding 21-hydroxylase) and *CYP11B1* (encoding 11β-hydroxylase type 1) was seen (Fig. 3e). Single-cell gene co-expression analysis revealed a distinct subset of cells that proportionately co-expressed *HSD3B2* and *CYP21A2* in early developmental stages, although by 19wpc it appeared that *CYP21A2* expression occurred in a greater number of cells, and that expression of *HSD3B2* (and its protein) was the likely rate-limiting factor (Fig. 3f).

Expression of *CYP21A2* and *CYP11B1* was linked (Supplementary Fig. 10). Taken together, these data suggest that there is an increase in gene expression of the enzymatic machinery needed for glucocorticoid synthesis across time.

It is also debated at what stage the developing fetal adrenal gland can synthesize mineralocorticoids, such as aldosterone^38, 39^. Here, *CYP11B2* (encoding 11β-hydroxylase type 2/aldosterone synthase) is a key enzyme in the final stages of aldosterone synthesis, as well as *HSD3B2*, which is needed to allow precursors into this pathway (Fig. 3d). In our scRNA- seq data, *CYP11B2* expression was low in early stages but increased by 19wpc in a sub- population of cells in the DZ (Fig. 3d, e). Again, single-cell co-expression analysis suggested that *HSD3B2* and *CYP11B2* are often linked (Fig. 3g) and their encoded enzymes are likely to be rate-limiting factors compared to those encoded by *CYP21A2* and *CYP11B1*.

### Transcriptional regulation of the fetal adrenal cortex

In order to study “core” transcriptional regulators of human adrenal cortex, we first identified genes that were differentially-expressed in the cortex cluster compared to non-cortex clusters at each scRNA-seq stage (log2FC>0.25, padj<0.05), and compared these genes to the Animal Transcription Factor Database (TFDB)^48^ (Supplementary Data 4). At each developmental time point studied, transcription factors represented between 1.8-2.4% of all differentially expressed cortex genes (Supplementary Fig. 11a). By intersecting these analyses, 17 “core” transcriptional regulators were identified that were common to all datasets (Fig. 4a-c, Supplementary Fig. 11, Supplementary Fig. 12). These factors were all present in bulk RNA-seq analysis of adrenal gland samples compared to control tissues (log2FC>1.5, padj<0.05), suggesting that they had a strong degree of adrenal specificity (Supplementary Fig. 11b, c; Supplementary Data 4).

**Fig. 4.**
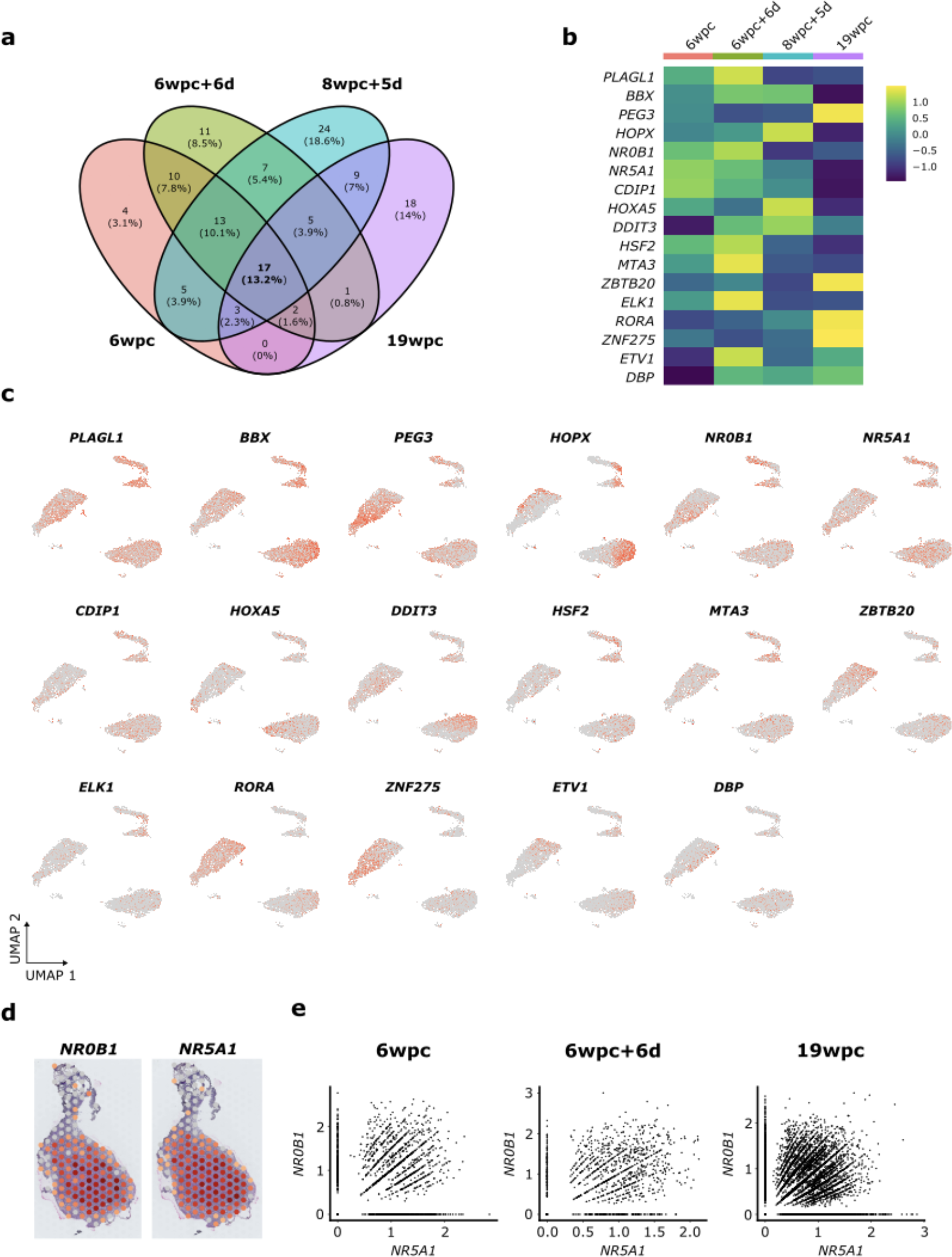
Expression of transcription factors during human adrenal cortex development. **a** Venn diagram showing the overlap of differentially-expressed transcription factors in the scRNA-seq dataset at each age. Differential expression was defined as being enriched in the adrenal cortex cluster compared to all other clusters in the whole adrenal sample at each age (log2FC>0.25, padj<0.05). A core group of 17 transcription factors common to all ages was identified. **b** Heatmap showing relative expression of these 17 transcription factors at each age in the scRNA-seq dataset. **c** Feature plots showing expression of these 17 transcription factors in adrenal cortex clusters (for annotation, see Fig. 3c). **d** Spatial transcriptomic spotplot expression of the key nuclear receptors, *NR0B1* (also known as DAX-1) and *NR5A1* (also known as steroidogenic factor-1, SF-1) at 7wpc+4d. **e** Scatter plots of expression of *NR5A1* in individual adrenal cortex single cells (scRNA-seq) compared to *NR0B1* (6wpc, 6wpc+5d, 19wpc).

Two key transcription factors that are well-established regulators of adrenal development are the orphan nuclear receptors, NR0B1 (DAX-1) and NR5A1 (SF-1)^15, 20, 22, 23^. Disruption of NR0B1 causes X-linked adrenal hypoplasia, which is one of the most common causes of PAI in children (boys)^13, 49^. NR5A1 is a master-regulator of adrenal and reproductive development and function, and more severe disruption is also associated with PAI in humans^23, 50^. Many studies have suggested that NR0B1 and NR5A1 can be functional partners, but data about expression in human development are still limited^15, 20, 33, 51^. Cluster analysis in scRNA-seq datasets as well as spatial transcriptomic analysis showed that expression of *NR0B1* and *NR5A1* occurs extensively throughout the fetal adrenal gland and that co-expression occurs in a subset of cells (Fig. 4c-e, Supplementary Fig. 11c). Taken together, these data support the key role that NR0B1 and NR5A1 play in transcription regulation and specification of human adrenal development.

### HOPX is a novel definitive zone factor

Although most of the “core” transcription factors identified showed expression throughout the adrenal cortex (i.e., DZ and FZ), one adrenal-enriched gene that was expressed very strongly in the DZ compared to the FZ was *HOPX* (Fig. 5a, b). HOPX is an atypical homeodomain protein (also known as Hop homeobox/“homeobox-only protein’) that lacks direct DNA-binding capacity but interacts with transcriptional regulators to maintain quiescence in specific embryonic and adult stem cell populations, and to control cell proliferation during organogenesis^52, 53^. HOPX also acts as a tumor suppressor, and reduced HOPX expression is associated with several cancers^52, 53^.

**Fig. 5.**
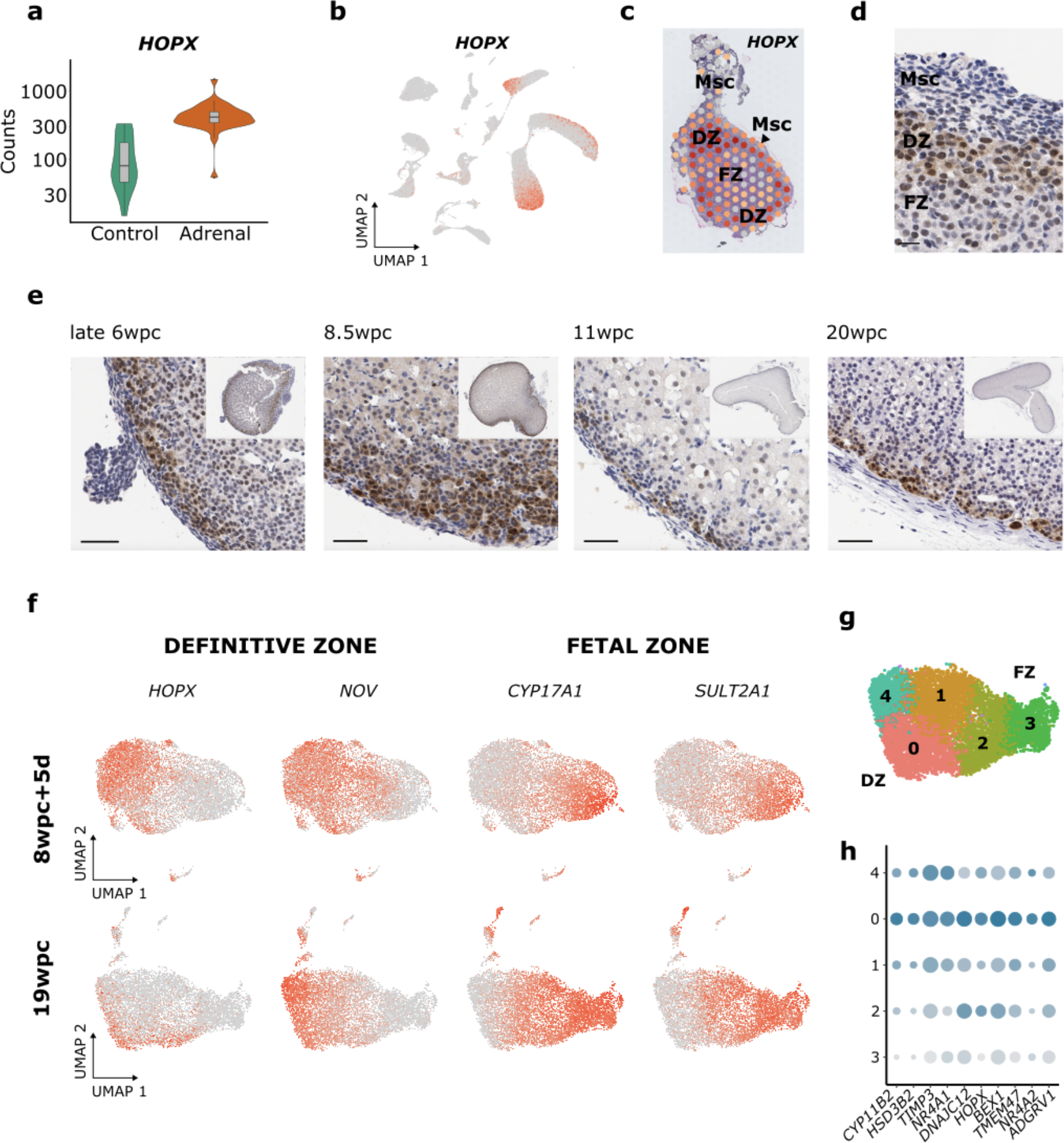
HOPX is a novel definitive zone factor. **a** *HOPX* expression (normalized counts) in the human developing adrenal gland (combined adrenal gland samples, bulk RNA-seq) compared to controls. **b** Feature plot of *HOPX* expression in the adrenal cortex clusters (for annotation, see Fig. 1g). **c** Spatial transcriptomic spotplot showing definitive zone (DZ) expression of *HOPX* at 7wpc+4d.FZ, fetal zone; Msc, mesenchyme. **d** Immunohistochemistry showing expression of *HOPX* in the DZ at late 6wpc between the layer of outer mesenchyme (Msc) and inner adrenal fetal zone (FZ) (scale 20μm). **e** Immunohistochemistry showing representative DZ expression of HOPX at each stage, with the whole adrenal gland inset. Scales all 50μm. **f** Feature plots of *HOPX* expression in the adrenal cortex cluster at two different ages (8wpc+5d, 19wpc) compared to the DZ marker *NOV*, and FZ markers *CYP17A1* and *SULT2A1*. **g** UMAP of key cortex clusters at 19wpc. **h** Dot-plot of the top differentially- expressed genes in cluster 0 compared to other clusters at 19wpc.

In our scRNA-seq dataset, HOPX was consistently one of the most differentially-expressed markers of the DZ compared to the FZ in all ages studied (Supplementary Fig. 13; Supplementary Data 3). This highly-specific enrichment of *HOPX* in the DZ was confirmed by spatial transcriptomic analysis, which showed a strong “ring” of *HOPX* DZ expression at 7wpc+4d, with a peripheral ring of weaker expression likely representing mesenchymal cells (Fig. 5c). This finding was validated by immunohistochemistry, which showed that HOPX defined the outer border of the DZ at the interface of the peripheral mesenchyme at late 6wpc (Fig. 5d). Furthermore, serial IHC analyses showed that HOPX was expressed in the outer DZ across time (late 6wpc-20wpc), marking the boundary between the developing adrenal gland and the mesenchyme (early) or subcapsular region of cells (later) (Fig. 5e, Supplementary Fig. 14).

As expected given its role in the DZ, *HOPX* co-localized in clusters with the DZ marker *NOV* in scRNA-seq analysis, especially during early stages of development (Fig. 5f). However, by 19wpc, *HOPX* expression was relatively reduced (Fig.4b, Fig. 5f) and localized within a zona glomerulosa-like cluster that also expressed *HSD3B2*, *CYP11B2*, and the orphan nuclear receptors *NR4A1* (NURR77)/*NR4A2* (NUR1) (Fig. 5g, h, Supplementary Fig. 13, Supplementary Fig. 15). Of note, an emergent population of NOV positive/HOPX negative cells was identified by scRNA seq at 19wpc, which was located just central to the peripheral HOPX positive cells on dual-labeled IHC (Fig. 5f, Supplementary Fig. 15e). Furthermore, *HOPX* does not seems to be expressed in the mature adult human adrenal gland, consistent with the decreased expression of this gene seen with time (https://www.proteinatlas.org/ENSG00000171476-HOPX/tissue). Thus, HOPX likely plays a role in defining the human fetal adrenal DZ and emerging zona glomerulosa in early development, and may maintain a specific population of cells in a replication state.

### Mesenchyme-cortex interactions during development

As the adrenal gland forms within a region of mesoderm/mesenchyme (Fig. 5c-e), more detailed analyses of potential ligand-receptor signaling interactions were undertaken using a combined adrenal cortex-mesenchyme scRNA-seq data. Notably, a potential transcriptomic “bridge” between the mesenchyme and cortex was identified in the merged adrenal dataset, particularly in the 6wpc+5d sample (Fig. 1g, Fig. 6a). A trajectory of cells undergoing differentiation from the mesenchyme to cortex was also observed (Fig. 6b).

**Fig. 6.**
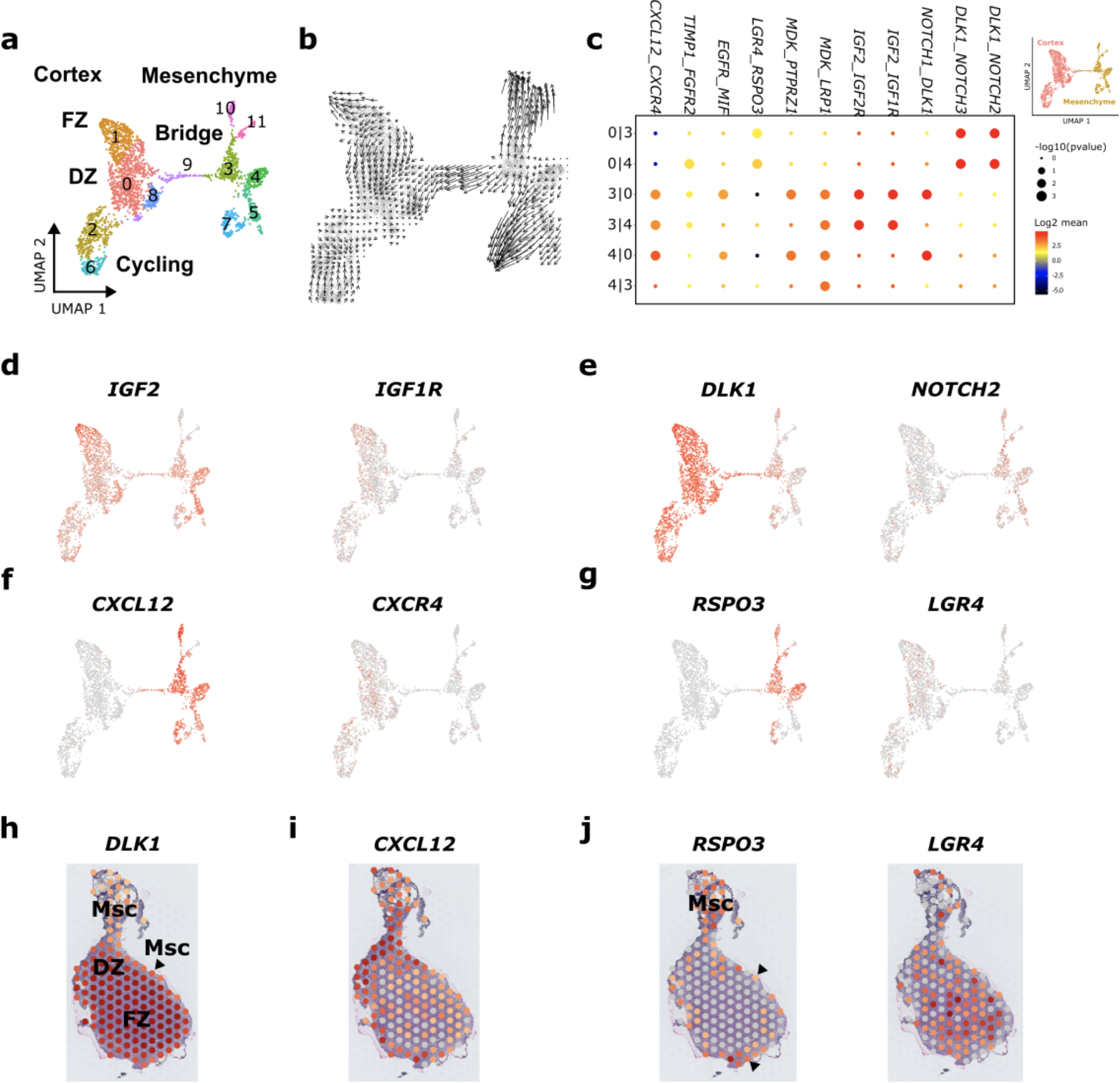
Potential bidirectional signaling interactions between the mesenchyme cluster and adrenal cortex. Data (scRNA-seq) shown at 6wpc+6d. **a** UMAP of the mesenchyme and adrenal cortex clusters demonstrating the potential “bridge” between the two populations of cells. Subclusters used for cell-cell communication analysis are shown. DZ, definitive zone; FZ, fetal zone. **b** Single-cell velocity estimates overlaid on the UMAP projection of mesenchyme-adrenal cortex clusters (RNA Velocity). **c** Potential ligand-receptor interactions for key subclusters in the mesenchyme (3, 4) and adrenal cortex (0), using CellPhoneDB. **d** Feature plot showing expression of *IGF2* (encoding ligand) and expression of *IGF1R* (encoding cognate receptor). **e** Feature plot showing expression of *DLK1* (encoding ligand) and expression of *Notch 2* (encoding receptor) (see also Supplementary Figure XX). **f** Feature plot showing expression of *CXCL12* (encoding ligand) and expression of *CXCR4* (encoding receptor). **g** Feature plot showing expression of *RSPO3* (encoding ligand) and expression of *LGR4* (encoding receptor). **h** Spatial transcriptomic spotplot (7wpc+4d) of *DLK1* in the definitive zone (DZ) and fetal zone (FZ) of the adrenal gland, with weaker expression in the mesenchyme (Msc)/subcapsular region. **i** Spatial transcriptomic spotplot (7wpc+4d) of *CXCL12*, strongest in the mesenchyme (Msc)/subscapsular region of the adrenal gland. **j** Spatial transcriptomic spotplot (7wpc+4d) of *RSPO3* in the mesenchyme (Msc, and arrowheads)/subcapsular region of the adrenal gland and *LGR4* in the adrenal cortex.

Using CellPhoneDB (CellPhoneDB v2.0)^54^ to investigate cell-cell communication networks and ligand-receptor interactions at this stage of adrenal gland development, several key systems were found to be enriched (Fig. 6c-g). For example, insulin-like growth-factor 2 (*IGF2*) showed strong expression in mesenchyme and adrenal cortex, with strongest expression in the fetal zone, the region of highest expression of its cognate receptors, *IGF1R* and *IGF2R* (Fig. 6c, d, Supplementary Fig. 16, Supplementary Fig. 17). In contrast, *DLK1* (also known as PREF1) showed high adrenal cortex expression whereas the linked Notch family of receptors are expressed predominantly in the mesenchymal component (Fig. 6c, e, h, Supplementary Fig. 16).

**Fig. 7.**
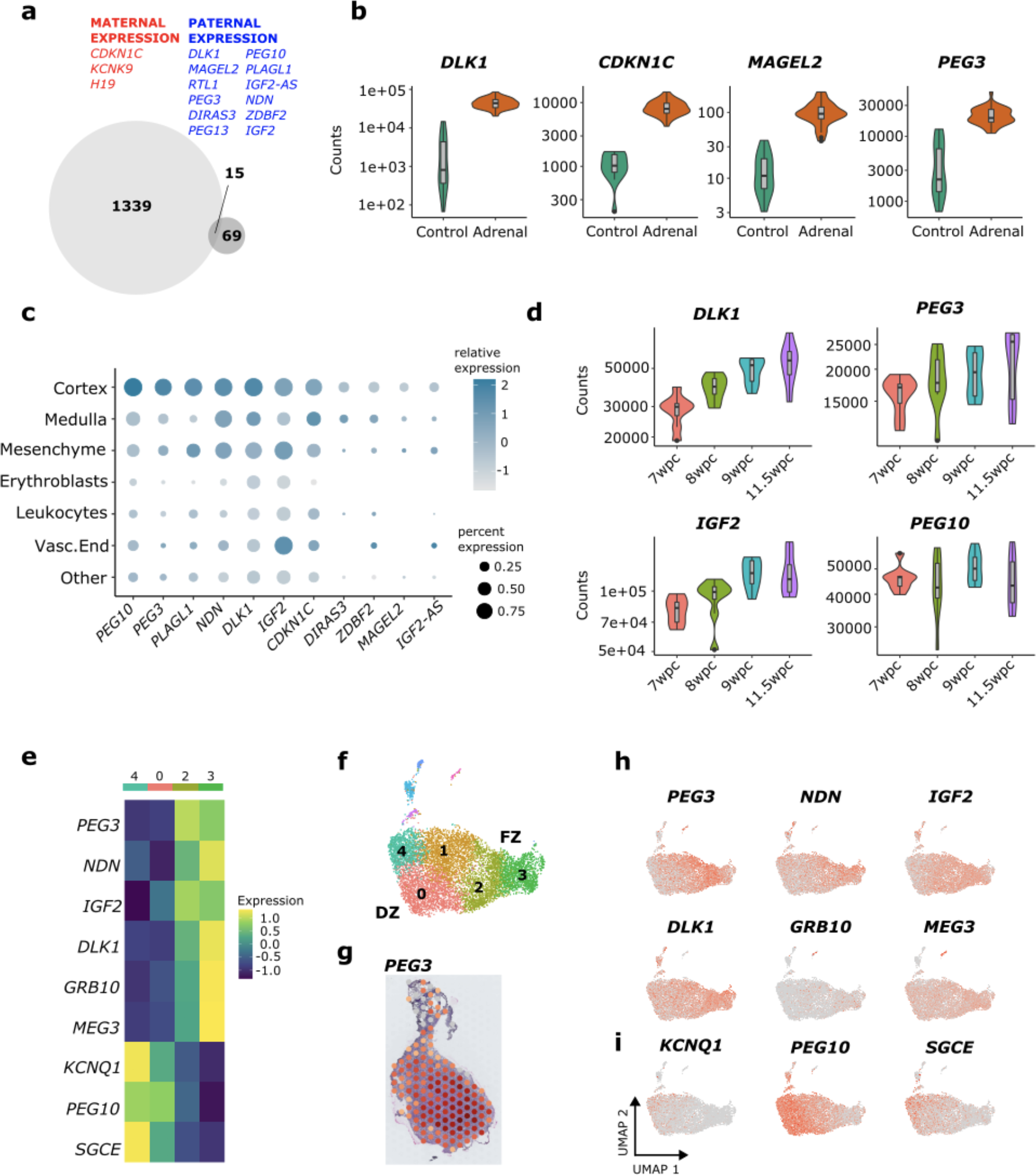
Imprinted genes in human adrenal development. **a** Venn diagram showing the 15 imprinted genes (non-placental specific) that are differentially-expressed in the adrenal cortex cluster (bulk RNA-seq adrenal > control, log2FC>1.5 padj<0.05). **b** Violin plots (normalized counts) of bulk RNA-seq expression of several key imprinted genes in the adrenal gland compared to control tissues. **c** Dot plot of key differentially-expressed imprinted genes in different scRNA-seq clusters of the developing human adrenal gland. **d** Violin plots showing time-series bulk RNA-seq expression of key imprinted factors in the developing human adrenal gland. **e** Heatmap of the expression of key imprinted genes in different clusters of the adrenal cortex at 19wpc (see Fig. 6f). **f** UMAP of adrenal cortex subclusters at 19wpc. **g** Spatial transcriptomic spotplot (7wpc+4d) of paternally-expressed gene 3 (*PEG3*) showing strong expression, especially in the central fetal zone. **h** Feature plots of seven key paternally-expressed (maternally-imprinted) genes in the adrenal cortex (19wpc). **i** Feature plot of three key maternally-expressed (paternally-imprinted) genes in the adrenal cortex (19wpc).

Two key signaling systems where ligands are potentially secreted from mesenchymal cells and have receptors in the developing adrenal cortex are *CXCL12* (encoding the ligand)/*CXCR4* (encoding the receptor), and *RSPO3* (ligand)/*LGR4* (receptor) (Fig. 6 c, f, g, i, j, Supplementary Fig. 16). *RSPO3*/*LGR4* are part of the WNT signaling system and *Rspo3*/Rspondin3 has been proposed previously to be a key ligand released by subcapsular cells in both mouse and human adrenal development, with potential interactions with Lgr5 and Znrf3^18, 55, 56^. Using spatial transcriptomics, *RSPO3* expression was found to be expressed in the mesenchyme, including in an outer layer around the early adrenal gland (7wpc+4d), whereas *LGR4* was expressed more centrally in the fetal zone region (Fig. 7g, j). Strong LGR5 and LGR6 expression or interactions were not seen (Supplementary Fig. 16). Thus, although several signalling systems have been proposed in adrenal development from data in the mouse^25, 27, 57^, our unsupervised analysis of ligand-receptor interactions support the roles of *IGF2*, *DLK1* and *RSPO3*/Rspondin3 as major components in human adrenal development, and suggest that *CXCL12* may also influence potential mesenchyme-adrenal interactions.

### Imprinted genes are enriched in the human fetal adrenal gland

*IGF2* and *DLK* are both imprinted genes, and it is well recognized that imprinted genes play a key role in many aspects of fetal and placental growth in humans^58^. Paternally-expressed (maternally-imprinted) genes are frequently linked to growth promotion, whereas maternally-expressed (paternally-imprinted) genes are associated with growth restriction.

To address the potential role of imprinted genes in the developing fetal adrenal gland in more detail, differential expression was initially studied using bulk-RNAseq data (adrenal versus control, log2FC>1.5 padj<0.05). We found that 15 out of 84 (17.9%) well-established, non-placental-specific human imprinted genes^59^ were differentially expressed in the adrenal gland, representing a significant enrichment of imprinted genes (15/1354 versus 69/18325, Chi-sq 15.9, p<0.0001) (Fig. 7a, b; Supplementary Data 5). Expression of these genes in adrenal cortex clusters was confirmed by scRNA-seq analysis (Fig. 7c). Several key paternally-expressed genes were identified (e.g. *DLK1*, *PEG3*, *IGF2*, *PEG10*), often in the FZ (Fig. 7 e-i). Taken together, these data highlight the important growth-promoting role paternally-expressed genes such as *IGF2* and *PEG3* play in the human fetal adrenal cortex during early development, at a time of rapid adrenal gland growth (Fig. 1b, Supplementary Fig. 17).

### Adult adrenal transcriptomic expression and primary adrenal insufficiency (PAI)

Finally, we considered how the transcriptomic profile of adrenal gland development relates to the adult adrenal gland, as well as to genes known to cause PAI.

Using the top differentially-expressed genes in the adult adrenal gland (n=12) (Human Protein Atlas, https://www.proteinatlas.org/humanproteome/tissue/adrenal+gland), we found consistent correlations with many differentially-expressed genes during developmental (Fig. 8a), although *GML* (encoding glycosylphosphatidylinositol anchored molecule like) was not present in the fetal data and several other genes were predominantly expressed in later fetal adrenal stage (19wpc) (e.g. *HSD3B2*, *CYP11B2*, *ADGRV1)* (Supplementary Fig. 18). This finding contrasts with *HOPX*, which is predominantly expressed in the fetal adrenal but not in the adult organ.

**Fig. 8.**
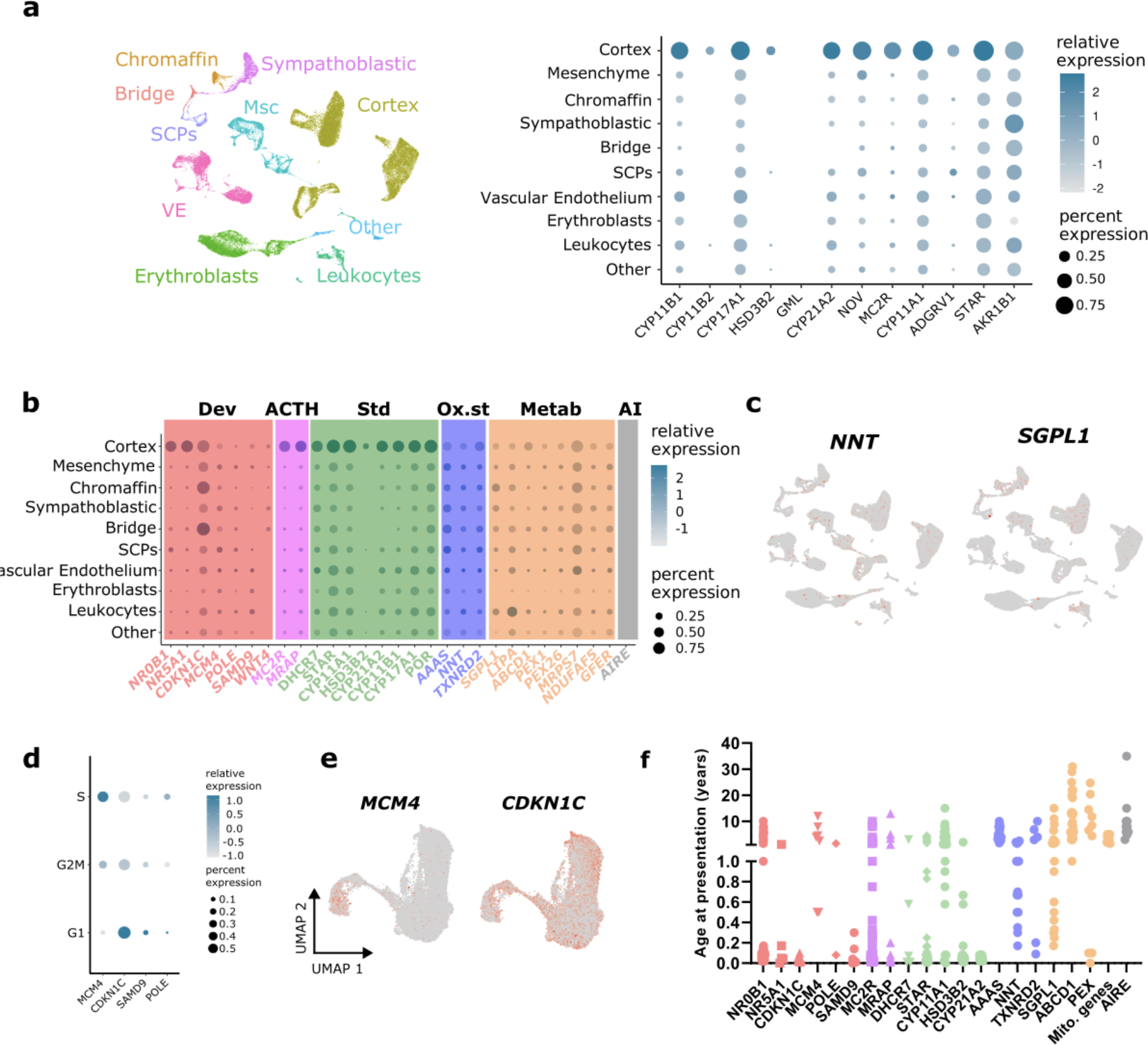
Expression of genes enriched in the adult adrenal gland and in monogenic causes of primary adrenal insufficiency. **a** Dot plot showing fetal adrenal gland expression of genes with the highest “tissue specificity score” (enriched expression) in the adult adrenal gland, as defined in the Human Protein Atlas (www.proteinatlas.org). **b** Dot plot showing the expression of genes associated with monogenic causes of primary adrenal (glucocorticoid) insufficiency (PAI) in the adrenal cortex and other adrenal clusters during development (see UMAP Fig. 8a). **c** Feature plot for expression of nicotinamide nucleotide transhydrogenase (*NNT*) and sphingosine-1-phosphate lyase 1 (*SGPL1*). **d** Dot plot of the expression of PAI- causing genes proposed to be involved in adrenal growth and cell division in different cell cycle phases (S phase, G2M, G1). **e** Expression of mini-chromosome maintenance complex component 4 (*MCM4*) and cyclin-dependent kinase inhibitor 1C (*CDKN1C*) in the integrated adrenal cortex cluster with cycling cells included (see Fig. 2i). **f** Age at presentation with adrenal insufficiency of children and young people with selected monogenic causes of PAI.

We also analyzed developmental expression of genes known to be monogenic causes of PAI (Fig. 8b)^10^. Most key transcription factors (e.g. *NR5A1*, *NR0B1*), components of steroidogenesis (e.g. *STAR*, *CYP11A1*, *CYP21A2*) and genes involved in ACTH-signalling (e.g., *MC2R*, *MRAP*) showed high specificity for expression in the fetal adrenal cortex cluster (Fig. 8b). However, many genes linked to oxidative stress processes or metabolic function were more ubiquitously expressed (e.g., *NNT*, *AAAS*, *SGPL1, ABCD1*)^60–63^ (Fig. 8b, c). In addition, out of those genes associated with multisystem growth restriction phenotypes (e.g. *MCM4*, *CDKN1C*, *SAMD9*, *POLE*)^64–66^, only *MCM4* (PAI, short stature, natural-killer cell deficiency) was expressed predominantly in cycling cells (S-phase) (Fig. 8b, d). Overall, these findings could have clinical relevance. For example, when the age of presentation of children with classic monogenic causes of PAI was analyzed, it emerged that children who had disruption of highly adrenal cortex/adrenal specific genes often presented with adrenal insufficiency soon after birth (in the first two weeks), whereas those children with defects in genes with less adrenal cortex specific profiles often have a delayed clinical presentation in later infancy, childhood or even adult life (Chi-sq 8.56, p<0.005) (Fig. 8e, Supplementary Data 6). These data suggest that, for some conditions, a period of postnatal stress and decompensation may be required before adrenal insufficiency presents, which could provide a window of opportunity for intervention if a diagnosis is made early enough.

## Discussion

This study provides one of the first detailed insights into the complexities of human adrenal gland development up to 20 wpc and demonstrates the benefits of integrating transcriptomic data (bulk RNA-seq, scRNA-seq, spatial) with developmental anatomy and physiology when investigating the biological basis of organogenesis and related clinical conditions.

It is already established that the human adrenal gland undergoes marked growth throughout gestation, and at birth is approximately the same weight as in adult life^37, 38^. Much of this growth is due to the expansion of the large FZ, which is only found in humans and higher primates. Here, we document changes in growth and morphology up to 20 wpc. Using scRNA-seq analysis of cycling cells, coupled with IHC markers of cell division (KI-67), we show that there is rapid cell division during the late embryonic/early fetal stage, and that the majority of dividing cells are located in the outer DZ region. A potential trajectory of cell differentiation from the DZ to FZ was seen during early adrenal development^25, 27, 43–46^.

Imprinted genes, such as *IGF2* and *DLK1*, play a key role in adrenal growth^36, 58, 67–70^. Here, we demonstrate strong expression of paternally-expressed growth-promoting genes, especially in the FZ region, consistent with the rapid growth seen during this stage of development.

Another key finding was the marked increase in vascularization of the adrenal gland across this time frame, and development of vascular sinusoids within the FZ. Novel imaging techniques such as microCT^71^ highlighted the surface arrangement of these vascular networks, especially on the inferior aspect of the adrenal gland that is adjacent on the upper pole of the kidney. Studies of angiogenesis and vascular remodeling in the fetal adrenal gland have focussed on both the VEGF/VEGFR1 and angiopoietin/Tie systems^72–75^. Vascular channels are crucial for transporting large amounts of adrenal androgens into the fetal circulation, with subsequent placental conversation to estrogens. The vascular anatomy also influences the dynamic interplay between the adrenal cortex and medulla, and children with steroidogenic defects in the cortex such as CAH have reduced medullary reserve^76–78^. The late embryonic and fetal period is a key time when these vascular systems are established.

Although the main role of the adult adrenal cortex is the biosynthesis and release of steroid hormones (mineralocorticoids, glucocorticoids, androgens), the extent to which these hormones can be generated in the fetal adrenal gland remains to be fully elucidated. Recent studies have looked at expression of key components of these pathways, or attempted to measure the major steroid hormones and their metabolites directly^38, 39^. Here, we show that the FZ has the transcriptomic machinery to secrete large amounts of adrenal androgens, such as DHEA(S). Precursors are shunted into this pathway due to the lack of HSD3B2 (encoding 3β-hydroxysteroid dehydrogenase type 2). Expression of the adrenocorticotropin (ACTH) receptor (*MC2R*) and its accessory protein (*MRAP*) increased with age, and showed strong expression in the FZ region. This finding is in keeping with ACTH-dependent stimulation of androgens in fetal adrenal cell or tissue cultures, suggesting the FZ androgen biosynthesis has the capacity to be ACTH-driven^79–81^.

In contrast, glucocorticoid biosynthesis (e.g. cortisol) requires *HSD3B2* expression. Consistent with two previous reports^37, 39^, we detected a potential transient increase in *HSD3B2* at around 8.5 wpc, but expression was more consistent by 19wpc in DZ cells that often co-expressed *CYP21A2* (encoding 21-hydroxylase) and *CYP11B1* (encoding 11 β- hydroxylase). Very limited expression of the genes required for mineralocorticoid biosynthesis (e.g. aldosterone) was seen early on, but a small proportion of DZ cells did express *CYP11B2* together with other relevant enzymes by 19wpc. This finding is consistent with a lack of aldosterone synthesis in the first half of gestation, although increases in CYP11B2 expression towards the end of the second trimester suggest the capacity for aldosterone synthesis is being established^38, 39^. Of note, pre-term babies often have hypotension and salt-loss, which may in part be due to immature development of mineralocorticoid biosynthesis, as well as relative mineralocorticoid resistance.

Understanding the dynamic transcriptomic and physiological changes around this time is key.

Two key transcription factors (TFs) that regulate fetal adrenal development are the nuclear receptors *NR0B1* (DAX-1) and *NR5A1* (SF-1)^15, 20, 22, 23, 82^. These genes encode two important nuclear receptors within a “core” set of 17 transcription factors identified, that were consistently differentially expressed in the adrenal cortex across time. Another transcription regulator identified that was remarkable for its consistent differential expression in the DZ compared to the FZ was *HOPX*^52^. HOPX is an atypical homeobox factor that lacks direct DNA binding and likely interacts with transcriptional regulators^52^, so is not universally classified as a TF. Nevertheless, HOPX is emerging as a key embryonic and adult stem cell marker involved in stem cell maintenance/quiescence^83, 84^ and controlled tissue differentiation^52^.

HOPX is reported to play a role in the development of mesoderm progenitor cells/hematopoietic stem cells^84, 85^, osteogenic cells, neuronal tissue^83^, cardiomyoblasts^86^, intestinal crypt/colonic cells^53, 87^, skin^88^, alveolar epithelial cells (Type I)^89^ and endometrium. HOPX can influence tissue repair and regeneration^87^, and reduced HOPX expression (through promoter methylation) is associated with several cancers (e.g. colon^53^, breast^90^, thyroid, pancreas^91^) and metastasis risk (e.g. nasopharyngeal^92^), suggesting HOPX acts as a tumor suppressor. Interactions with WNT signalling^86^, activated SMAD^86^ and CXCL12^84^ have also been proposed. Our findings support a recent report of HOPX expression in early human adrenal development^18^, but suggest a developmentally-important time course for HOPX into later gestation and postnatal life. We show clearly that HOPX is expressed at the outer boundary of the developing DZ, close to the mesenchymal layer initially and in the subcapsular part of the DZ later. Given the decrease of HOPX with age, it is possible that HOPX plays a role in maintaining controlled cell proliferation and growth in the developing DZ. Of note, Xing et al showed in 2010 that HOPX is downregulated following ACTH stimulation in studies of both adult and fetal adrenal cells *in vitro*, whereas ACTH stimulates synthesis of steroidogenic enzymes^79^. Coupled with our observation of strongest *MC2R*/*MRAP* expression in the FZ, we hypothesize that ACTH and the ACTH pathway may promote adrenal differentiation not only through upregulating steroidogenesis, but through downregulating *HOPX*/HOPX and allowing cells to actively differentiate. Certainly, the role for HOPX in defining the DZ during early development and differentiation warrants further investigation.

As the adrenal gland arises from a condensation of intermediate mesoderm/mesenchyme at around 4wpc, we also focussed on mesenchyme-cortex interactions during the earliest phase of development investigated (6wpc to 8wpc). Indeed, IHC and spatial transcriptomic analysis clearly showed the adrenal gland developing within an outer ring of mesenchymal cells next to a mesenchymal “pedicle”. The early adrenal gland had a bulk transcriptomic profile closer to the kidney (mesoderm) initially that became increasingly distinct with time, as the relative proportion of mesenchymal cells diminished and that of adrenal-specific cells increases. By studying mesenchymal-cortex clusters at 6wpc we identified a potential trajectory of cells differentiating from the mesenchyme to the cortex, consistent with a pool of progenitor cells in region, which ultimately locate within the subcapsular region^28, 93, 94^.

Several different signaling systems have been proposed to regulate mesenchyme-cortex interactions from studies in the mouse.^25–28, 80, 95^ Using an unsupervised approach of CellPhoneDB^54^ to identify ligand-receptor interactions, we found that Rspondin-3 (*RSPO3*) could have an important role. Rspondin-3 is a component of the WNT-signalling pathway and has been shown to be expressed in the subcapsular region of cells in the developing mouse adrenal gland^55^, as well as in subcapsular cells in the 8wpc human adrenal gland^18^, and potentially mediates a gradient of WNT signalling involved in adrenal zonation.

Although interactions with LGR5 have been suggested^55^, we identified LGR4 as the most likely expressed putative cortex receptor. A potential role for CXCL12 (mesenchymal ligand) and CXCR4 (adrenal cortex receptor) was also identified. Other signaling systems proposed from mouse models (eg *Shh*/*Gli*) were not found to be strongly expressed in the developing human adrenal gland at this stage. Taken together, these data suggest the Rspondin3-driven WNT signalling has a key role in human adrenal development, as well as in mice.

Our findings also address how basic biological mechanisms relate to human disease. Our translational focus over the years has been on monogenic causes of PAI. In children and young people these conditions are often inherited and represent potentially life-threatening disorders needing prompt diagnosis and management^9^. Progress over the past three decades has identified around 30 single gene causes of PAI^10^,some of which are shown here to have specific developmental features (e.g., NR5A1/NR0B1 as core transcriptional regulators^23^; MCM4 in S-phase cell division^96^). The novel differentially-expressed adrenal genes found in our analyses will provide candidate genes for new genetic causes of PAI in the future. Furthermore, the unexpected observation that several key genes associated with PAI are not differentially expressed, and the clinical conditions they cause generally present with PAI at a later age, suggests that a period of postnatal stress/decompensation is required for the adrenal insufficiency to manifest. Making an early diagnosis – potentially even through newborn genetic screening programmes – means a window of opportunity exists to alter the disease course, or at least to predict the onset of PAI and avoid an adrenal crisis.

Insights into basic mechanisms of human adrenal development also have implications for better understanding the drivers of adrenal tumors. We have previously shown the opposing effects of variants in *CDKN1C*/CDKN1C, whereby gain-of-function of this cell-cycle repressor is associated with adrenal hypoplasia and IMAGe syndrome, and loss of function is associated with Beckwith-Wiedemann syndrome with an adrenal neoplasm risk^65^. In childhood especially, adrenocortical tumorigenesis has been linked to increased expression of NR5A1^97–99^ and IGF2 (through aberrant regulation of the 11p H19/IGF2 imprinting locus)^99–101^, and IGF1R blockade has been explored as a treatment for adrenal tumors in experimental models and trials^102–104^. Thus, the association of imprinted genes (e.g., *CDKN1C*, *IGF2*) with growth and tumor risk is emerging. More recently, CXCR4 expression has been used as a marker and potential therapeutic target in adrenal cancer^105, 106^, as well as for clinical diagnostic imaging of aldosterone secreting adenomas using ^68^Ga-pentixafor PET/CT^107–110^. Our findings also have relevance for the mechanisms of adrenal androgen synthesis and regulation in CAH (e.g., 21-hydroxylase deficiency)^3, 40, 47^, for insights into adrenarche and links between the FZ and zona reticularis^5, 111, 112^, and for potential “programming” of the hypothalamic-pituitary-adrenal (HPA) axis during development, which could have implications for postnatal variability in HPA axis function and stress responses^113, 114^.

These data have several limitations. The developmental period of tissue accessibility was somewhat limited and a greater sample number over time would have provided more detailed data. Also, whilst scRNA-seq and spatial transcriptomic platforms provide significant new insight, the ability to obtain increased sequencing reads per cell, more cells sequenced per sample, or greater spatial resolution is always improving and will help address some of the hypotheses generated here in the future. Understanding anatomical and physiological relations during development will be key going forward, at gene transcription, RNA expression and protein levels, and integrating detailed histology and imaging with basic cell biology will be crucial, as we have attempted to do here.

In summary, this study highlights the unique developmental complexities of human fetal adrenal gland development up unto mid-gestation, and provides an integrated transcriptomic roadmap with potential long-term consequences for human health and disease.

## Methods

### Tissue samples

Human embryonic and fetal tissue samples used for bulk-RNA seq, immunohistochemistry and microCT were obtained with ethical approval (REC references: 08/H0712/34+5, 18/LO/0822, 08/H0906/21+5, 18/NE/0290) and informed consent from the Medical Research Council (MRC)/Wellcome Trust-funded Human Developmental Biology Resource (HDBR) (http://www.hdbr.org). HDBR is regulated by the U.K. Human Tissue Authority (HTA; www.hta.gov.uk) and operates in accordance with the relevant HTA Codes of Practice. The age of embryos up to 8wpc was calculated based on Carnegie staging, whereas in older fetuses the age was estimated from foot length and knee-heel length in relation to standard growth data. Samples were karyotyped by G-banding or quantitative PCR (chromosomes 13, 16, 18, 21, 22 and X and Y) to determine the sex of the embryo/fetus and to exclude any major chromosomal rearrangements or aneuploidies. The acquisition of adrenal samples used to generate scRNA-seq data and spatial transcriptomics has been described previously^24^, under the following studies: NHS National Research Ethics Service reference 96/085 (fetal tissues) and the joint MRC/Wellcome Trust–funded HDBR (as above). An overview of all samples used in the study is provided in Supplementary Data 1. Samples were stored in the appropriate media or at -80°C until processing. Adrenal dimensions were measured to the nearest 0.5 mm, using a light microscope when necessary. Adrenal weights (single gland, 10% formalin) were measured on an analytical balance (Pioneer PX, Ohaus) after removal of surface liquid.

### Micro-focus computed tomography (micro-CT)

The 17wpc adrenal gland studied (10% formalin) was immersed in 1.25% potassium tri- iodide (I2KI) at room temperature for 48 hours, then rinsed, dried and wax embedded^115^. Once hardened, excess wax was trimmed in order to preserve tissue shape, to reduce dehydration and movement artefact, and to optimize contact with the X-ray beam source. Micro-CT scans were carried out using a Nikon XTH225 ST scanner (Nikon Metrology, Tring, UK) with the following settings: Tungsten target, X-ray energy 110 kV, current 60 µA (power 6.6 Watts), exposure time 1420 ms, one frame per 3141 projections, detector gain 24 dB, and scan duration of 75 minutes. Modified Feldkamp filtered back projection algorithms were used for reconstructions within proprietary software (CTPro3D; Nikon Metrology) and post-processed using VG StudioMAX (Volume Graphics GmbH, Heidelberg, Germany) to create the images at 4.77 µm isotropic voxel sizes.

### Bulk RNA-seq

Total RNA was extracted from human fetal adrenal samples (n = 32; Fig. 1a, Supplementary Fig. 1) and controls (n=14, Fig. 1a, Supplementary Fig. 1, balanced across the age range) using the AllPrep DNA/RNA Mini Kit (Qiagen). cDNA libraries were prepared using the KAPA mRNA HyperPrep Kit (Roche) and subsequently sequenced on a NextSeq 500 sequencer (paired-end 43 bp) (Illumina) in a single run to reduce potential batch effects. Fastq files were processed by FastQC and aligned to the human genome (Ensembl, GRCh 38.86) using STAR (2.5.2a)^116^. The matrix containing uniquely mapped read counts was generated using featureCounts^117^, part of the R package Rsubsead. Differential-expression analysis was performed using DESeq2^118^, using eight control samples instead of 14 where indicated to prevent duplication of specific tissue-types. Heatmaps for distances between samples and differentially expressed genes in adrenal vs. control samples were generated using the pheatmap library in R.

### Single-cell RNA-seq (scRNA-seq)

Detailed methods have been reported previously for the single cell sequencing of the samples used, including fetal adrenal single-cell dissociation, 10X Chromium processing (Chromium Single Cell 3’ kit) (10X Genomics), cDNA library preparation and sequencing (Illumina HiSeq 4000). A processed single cell matrix was generated as described before^24^ with minor modifications. Unless specified, cycling cells were discarded from the analysis. The R package Seurat (v4.0.2)^119^ was used for processing the single cell matrix. Briefly, the count matrix was normalized and 2000 highly variable genes chosen. After gene scaling, dimensionality reduction was performed using the first 75 principal components (PCs). The FindClusters and RunUMAP functions were used to identify clusters and to allow UMAP visualization. The clustree package in R^120^ was used to select the resolution parameter for clustering. Differentially-expressed genes between clusters were calculated using the FindAllMarkers or FindMarkers functions using the parameters ‘min.pct=0.25 and logfc.threshold=0.25’ (Wilcoxon Rank Sum test). Internal functions in Seurat (FeaturePlot, RidgePlot) were used to visualize marker expression. The FeatureScatter function was used to generate plots for pair of genes. The dittoSeq Bioconductor package^121^ was used to generate barplots, heatmaps and dotplots. RNA velocity on selected fetal adrenal samples was calculated using velocyto and plotted using the velocyto.R package in R as described before^24^. Adrenal cortex sample integration was performed using datasets normalized with SCT as described in vignettes (Seurat). Cell-cell communication by ligand-receptor interactions was calculated using CellPhoneDB v.2.0^122^.

### Histology/Immunohistochemistry (IHC)

Human embryonic/fetal adrenal glands at four different ages (“late 6wpc”, 8.5wpc, 11wpc, 20wpc) were fixed in 4% paraformaldehyde before being processed, embedded and sectioned for histological analysis and immunohistochemistry (IHC). Standard hematoxylin and eosin (H&E) staining was performed on 4µm sections to show key structural regions and vasculature. IHC was undertaken on 4µm sections using a Leica Bond-max automated platform (Leica Biosystems). In brief, sections first underwent antigen retrieval to unmask the epitope (Heat Induced Epitope Retrieval (HIER), Bond-max protocol F), endogenous activity was blocked with peroxidase using a Bond polymer refine kit (cat # DS9800), then incubation was undertaken with the relevant primary antibody for 1 hour. The following primary antibodies were used: VEGFR1 (Thermo Fisher PA1-21731, 1:100 dilution, HIER1 for 20 mins), KI67 (Leica Ready to use clone K2 PA0230, 1:100, HIER 2 for 20 mins), NOV (Sigma Aldrich HPA019864, 1:100, HIER1 for 20 mins), SULT2A1 (Sigma Aldrich HPA041487, 1:100, HIER2 for 20 mins) and HOPX (Sigma Aldrich HPA030180, 1:100, HIER2 for 20 mins). Next, the post-primary antibody was applied to the sections (Bond polymer refine kit) and horseradish peroxidase (HRP) labelled polymer, followed by 3, 3-diaminobenzidine (DAB) chromogen solution to precipitate the locus of antigen-antibody interactions (all Bond polymer refine kit). Sections were counterstained with hematoxylin, washed in deionized water, dehydrated in graded alcohols, cleared in two xylene changes and mounted for light microscopy. The stained slides were imaged on an Aperio CS2 Scanner (Leica Biosystems) at 40x objective. Analysis was undertaken with QuPath (v.0.2.3) (https://qupath.github.io) and Leica ImageScope (Leica Biosystems) software.

Dual-staining was performed with anti-HOPX (1:100 dilution) and anti-NOV (1:100 dilution) antibodies (as above) on 20wpc human fetal adrenal gland. Antigen retrieval was heat induced, HIER2 20 mins. Staining was performed sequentially on the Bondmax autostainer using anti-HOPX detected by brown chromogen (Bond polymer refine kit, cat # DS9800) and anti-NOV detected by red chromogen (Bond polymer Red kit, cat # DS9390).

### Spatial transcriptomic analysis

Spatial transcriptomic analysis of a single adrenal gland (7wpc+5d) was undertaken based on a standard 10X Genomics Visium protocol (10X Genomics). In brief, the fresh adrenal sample was snap frozen and embedded in OCT. Cryosections (10 µm) were cut and placed on Visium slides. Sections were fixed in cold methanol and stained with H&E to visualize the structures and tissue integrity, before permeabilization, reverse transcription and complementary DNA synthesis. Second-strand cDNA was liberated and libraries (Single-index) were generated by a PCR-based protocol. Libraries were sequenced on a HiSeq400 sequencer (Illumina).

Sequencing data were aligned to GRCh38 human reference genome using Space Ranger Software to quantify gene counts in spots.

### Adult adrenal gland gene enrichment

Data for the most highly differentially-expressed (enriched) adult adrenal gland genes was derived from the Human Protein Atlas, using the “tissue specificity score”. The tissue specificity score (TS) represents the fold-change between the expression level in adrenal gland and that in the tissue with the second-highest expression level (https://www.proteinatlas.org/humanproteome/tissue/adrenal+gland).

### Primary adrenal insufficiency (PAI): clinical presentation

Data for the clinical age of presentation of children and young people with genetic causes of PAI was obtained from original case-series reports of classic and non-classic conditions using PubMed (https://pubmed.ncbi.nlm.nih.gov/; accessed July 2022). Relevant literature sources are shown in Supplementary Data 6. Where PAI is a rare association of a condition, or where limited data are currently available, all published individual case reports were reviewed by two observers (J.A., F.B.). “Early”-onset PAI was defined as having at least one report of an infant presenting with adrenal insufficiency within the first two weeks of life, and “late”-onset PAI after this time. PAI-associated genes were termed “adrenal specific” if bulk RNA-seq data showed greater expression of that gene in the adrenal gland compared to controls (log2FC>2, padj<0.05), and if expression in the integrated adrenal cortex cluster was high (Supplementary Data 5).

### Statistical analysis

Statistical analysis for bulk- and single-cell RNA-seq data is described above within packages of differential expression analysis, with adjustments for multiple corrections. Chi-square analysis was performed GraphPad (Prism). In all analyses, a p-value or adjusted p-value less than 0.05 was taken as significant.

## Data Availability

### Data repository links

Single-cell RNA-sequencing data are deposited in the European Genome-phenome Archive (accession code EGAD00001008345).

Bulk RNA-sequencing data are deposited in ArrayExpress/Biostudies (accession number E-MTAB-12492).

## Supplementary Files

Supplementary Movie 1: Micro-CT of fetal adrenal gland at 17wpc

Supplementary Figures: Supplementary figures 1-18

Supplementary Data 1: Overview of samples included in the study (phenodata)

Supplementary Data 2: Bulk RNA-seq differentially expressed genes (adrenal vs controls)

Supplementary Data 3: Gene expression markers (single cell clusters)

Supplementary Data 4: Adrenal cortex - transcription factors

Supplementary Data 5: Adrenal cortex - imprinted genes

Supplementary Data 6: Clinical data (age of presentation) and related references

## Acknowledgements

This research was funded in whole, or in part, by the Wellcome Trust Grants 209328/Z/17/Z, 216362/Z/19/Z, 206194, 211276/Z/18/Z, 108413/A/15/D, and 223135/Z/21/Z. For the purpose of Open Access, the author has applied a CC-BY public copyright license to any Author Accepted Manuscript version arising from this submission. J.C.A. had additional research support from Great Ormond Street Hospital Children’s Charity (grant V2518). ICS is funded by the National Institute of Health Research (NIHR) (ICA-CDRF-2017-03-53 and NIHR302390). Human fetal material was provided by the Joint MRC/Wellcome Trust (Grant MR/R006237/1) Human Developmental Biology Resource (http://www.hdbr.org). Research at UCL Great Ormond Street Institute of Child Health is supported by the National Institute for Health Research, Great Ormond Street Hospital Biomedical Research Centre (grant IS-BRC-1215-20012). The views expressed are those of the authors and not necessarily those of the National Health Service, National Institute for Health Research, or Department of Health. The funders had no role in study design, data collection and analysis, decision to publish, or preparation of the manuscript. We also thank other members of the Human Developmental Biology Resource, UCL Genomics and Sanger Institute for their additional contributions to this work.

## Author contributions

Author contributions were as follows. Study conceptualization: IdV, SB, JCA; Methodology: IdV, MDY, ICS; Investigation: IdV, MDY, OKO, ICS, FB, BC, NM, TB, PN, KS, JPS, SMM, UCL Genomics, JCA ; Formal analysis: IdV, MDY, GD, ISC, EK, FB; Data curation: IdV, MDY, FB; Resources: HDBR; Project administration: JCA; Supervision: OJA, SB, JCA; Validation: IdV, MDY, JCA; Visualization: IdV, OKO, ICS, FB, JCA; Writing – original draft: IdV, JCA; Writing – review & editing: All authors; Funding acquisition: SB, JCA.

