## Supplementary Figs 1-18 for "A cell atlas of human adrenal cortex development and disease"

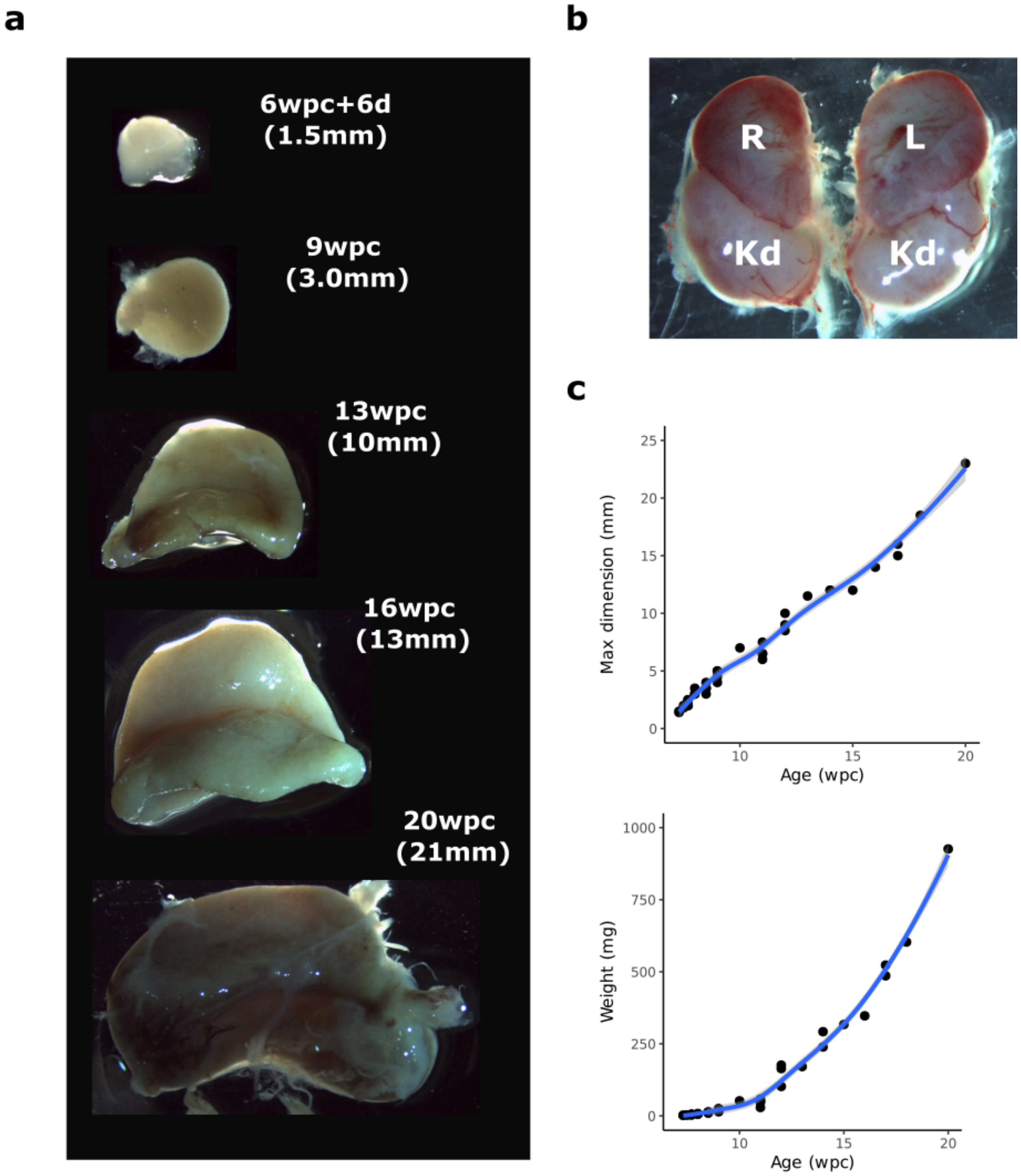


**Supplementary Fig. 1. Overview of human adrenal growth. a** Photographs of single adrenal glands at 6 weeks post conception (wpc)+6 days (d), 9wpc, 13wpc, 16wpc and 20wpc (not to scale). The corresponding maximum dimension is indicated. **b** Photograph of the right (R) and left (L) adrenal glands above the kidneys (Kd) at 14wpc. **c** Growth plots for adrenal size (maximum dimension) (*upper panel*) and weight (*lower panel*) between 7wpc and 20wpc (n=36).

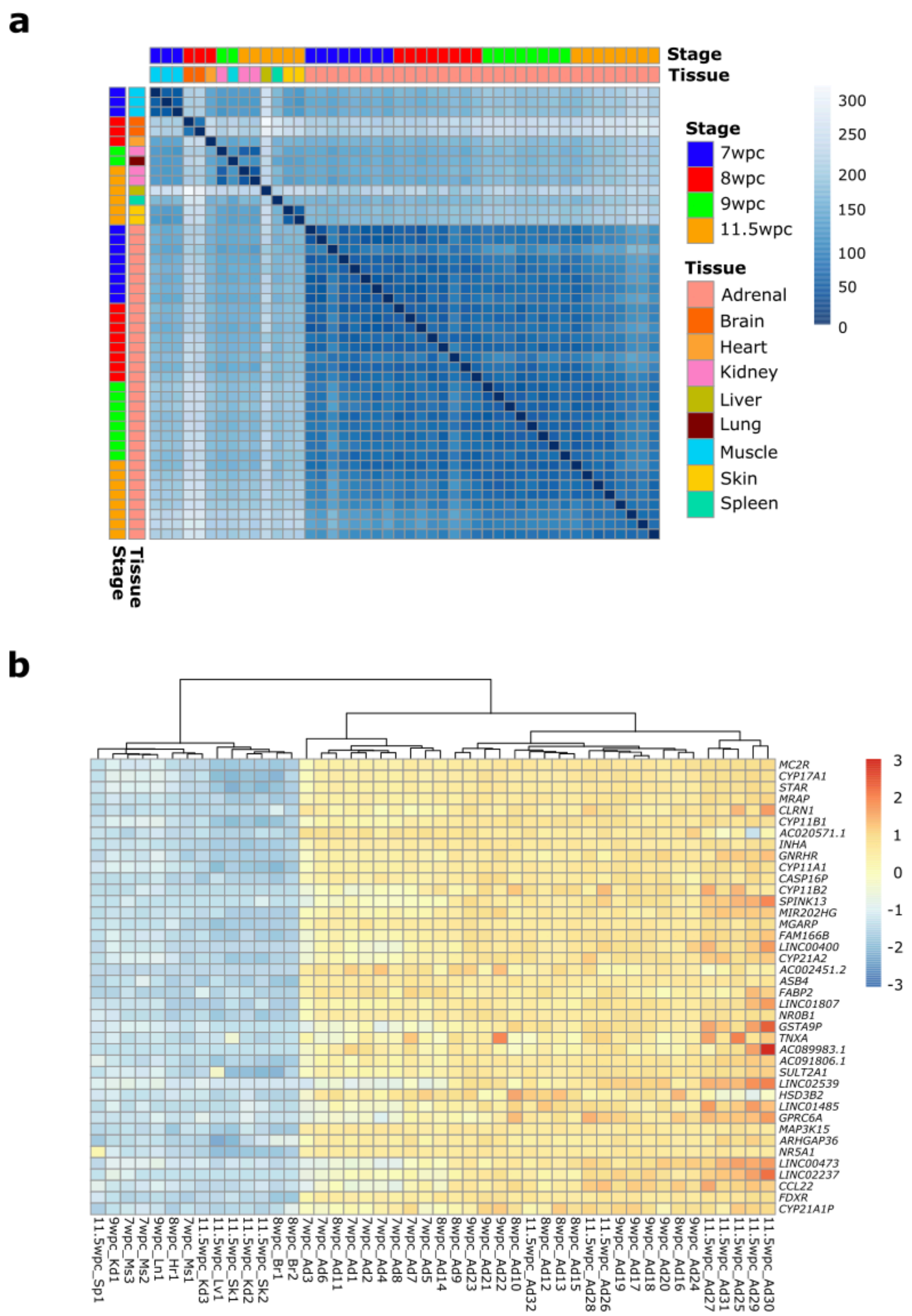


**Supplementary Fig. 2. Gene expression heatmaps across the bulk RNA-seq datasets. a** Correlation plot of gene expression in adrenal glands and controls across the time series. The origin and age of each tissue sample is shown. **b** Unsupervised clustering heatmap to show the top 50 differentially-expressed genes (based on fold change) in the adrenal samples compared to control tissues. Ad, adrenal; Br, brain; Hr, heart; Kd, kidney; Ln, lung; Lv, liver; Ms, muscle; Sk, skin; Sp, spleen; wpc, weeks post conception.

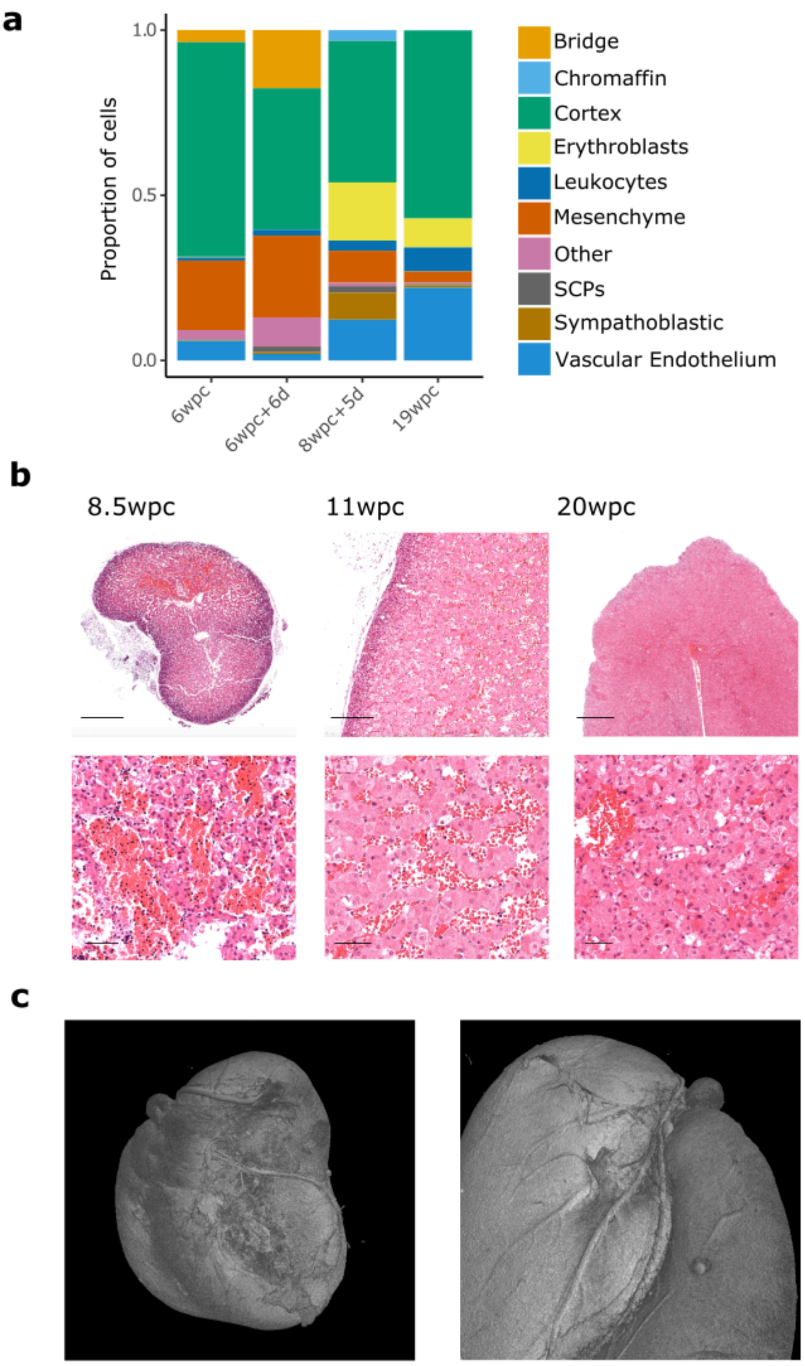


**Supplementary Fig. 3. Vascularization of the adrenal gland with development.** **a** Relative proportion of annotated clusters at each age. **b** Histology of adrenal glands to show the developing network of vascular sinusoids and central sinuses. Hematoxylin and eosin (H&E) staining. Scale bars: 8.5wpc, 400µm (upper), 50µm (lower); 11wpc, 200µm (upper), 50µm (lower); 20wpc, 800µm (upper), 50µm (lower). **c** MicroCT surface images showing vasculature at 17wpc.

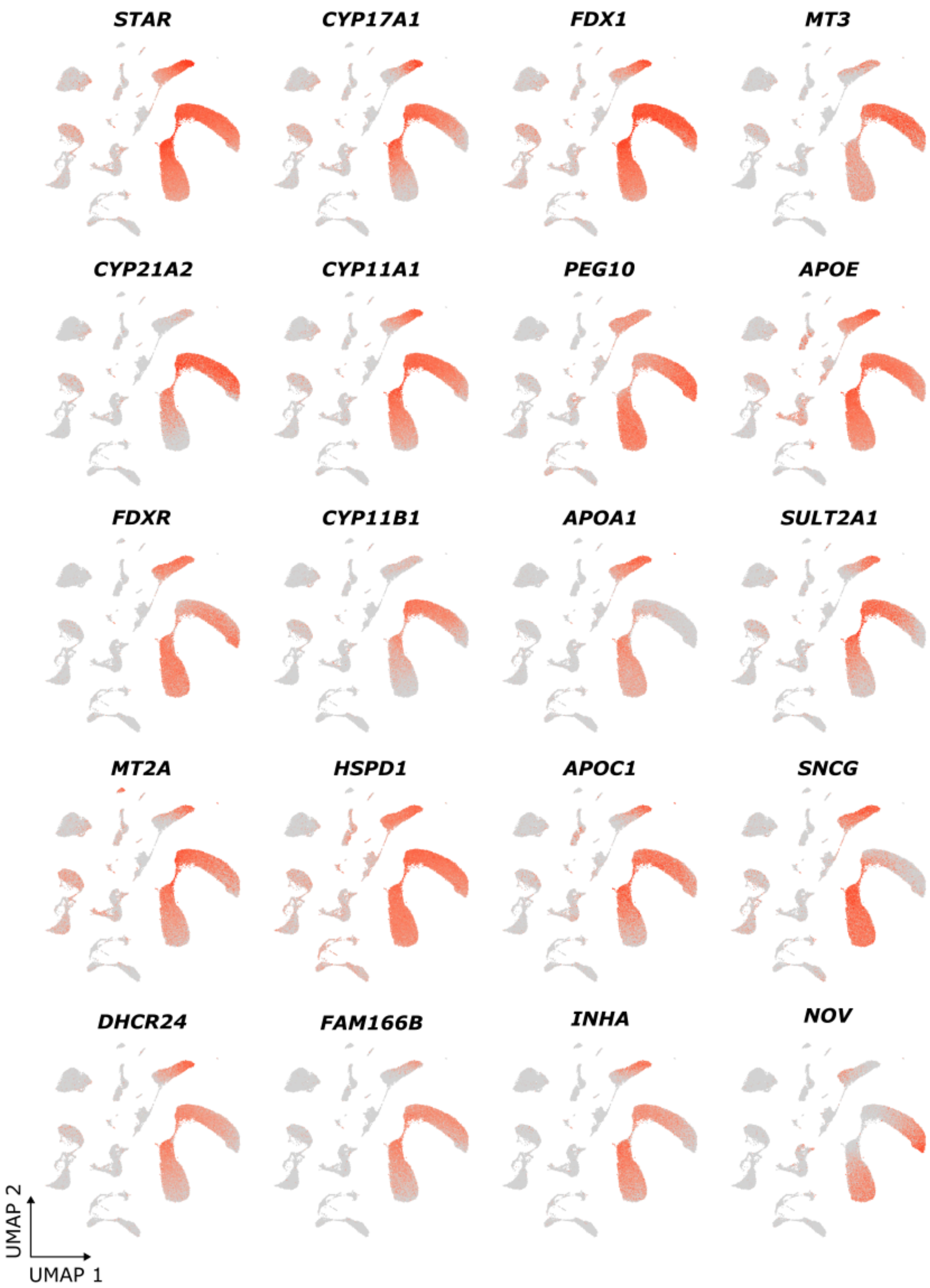


**Supplementary Fig. 4. Feature plots of the top differentially-expressed adrenal cortex cluster genes from the merged scRNA-seq dataset.** The annotated clusters of the UMAP are shown in main Fig. 1g and dot plot of expression of these genes is shown in main Fig. 2a.

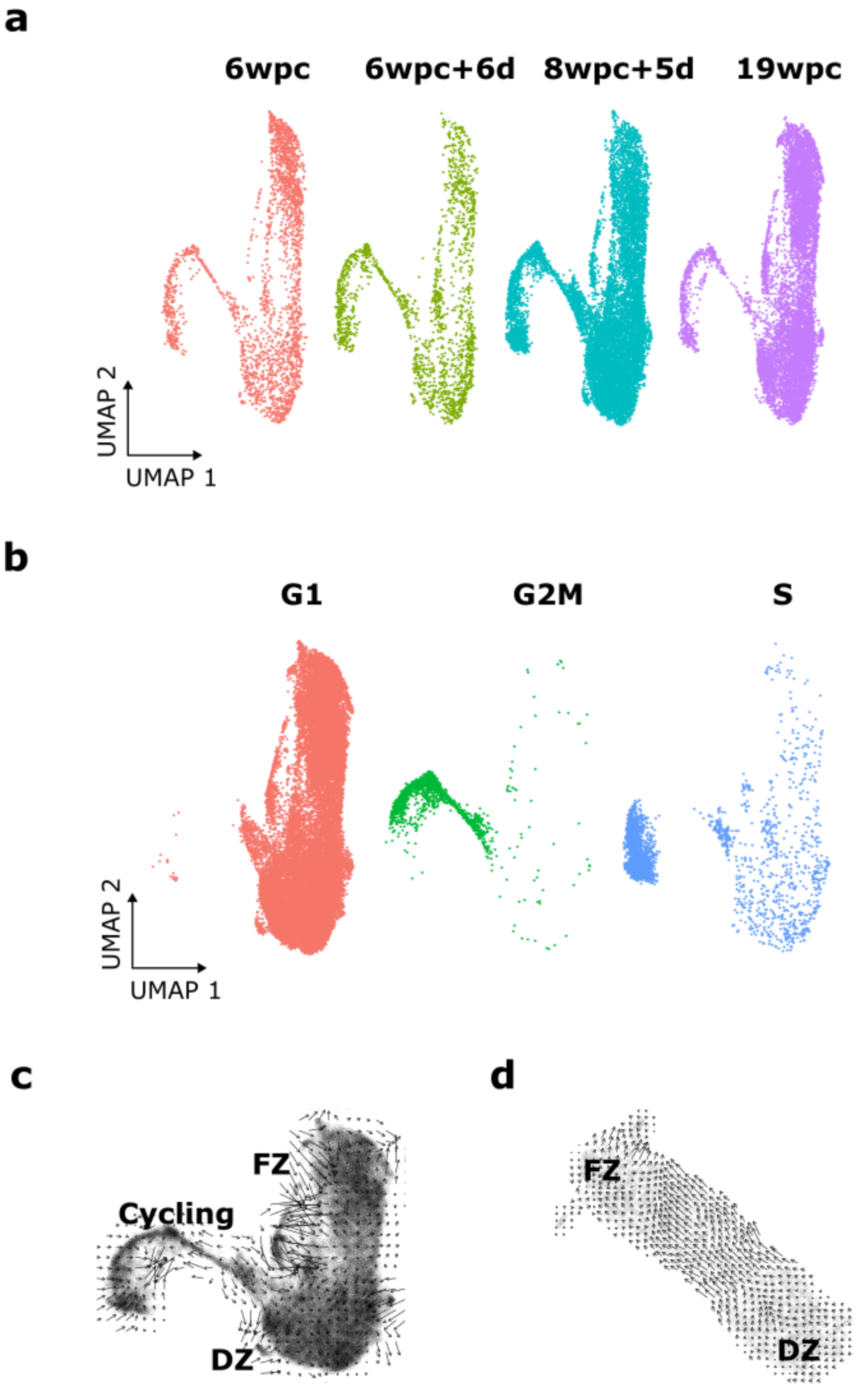


**Supplementary Fig. 5. Analysis of cycling cells and cell trajectories contributing to the integrated UMAP in main Fig. 2i. a** Overview of the cortex and cortex dividing cells at each age. **b** Integrated UMAP showing cells in G1 phase (red) and dividing cells in GM2 phase (green) and S phase (blue). **c** RNA velocity showing cycling cell trajectories from a node closest to the definitive zone (DZ) rather than the fetal zone (FZ). **d** RNA velocity of the 6wpc scRNA-seq adrenal cortex cluster showing a potential trajectory from the DZ to FZ at this age.

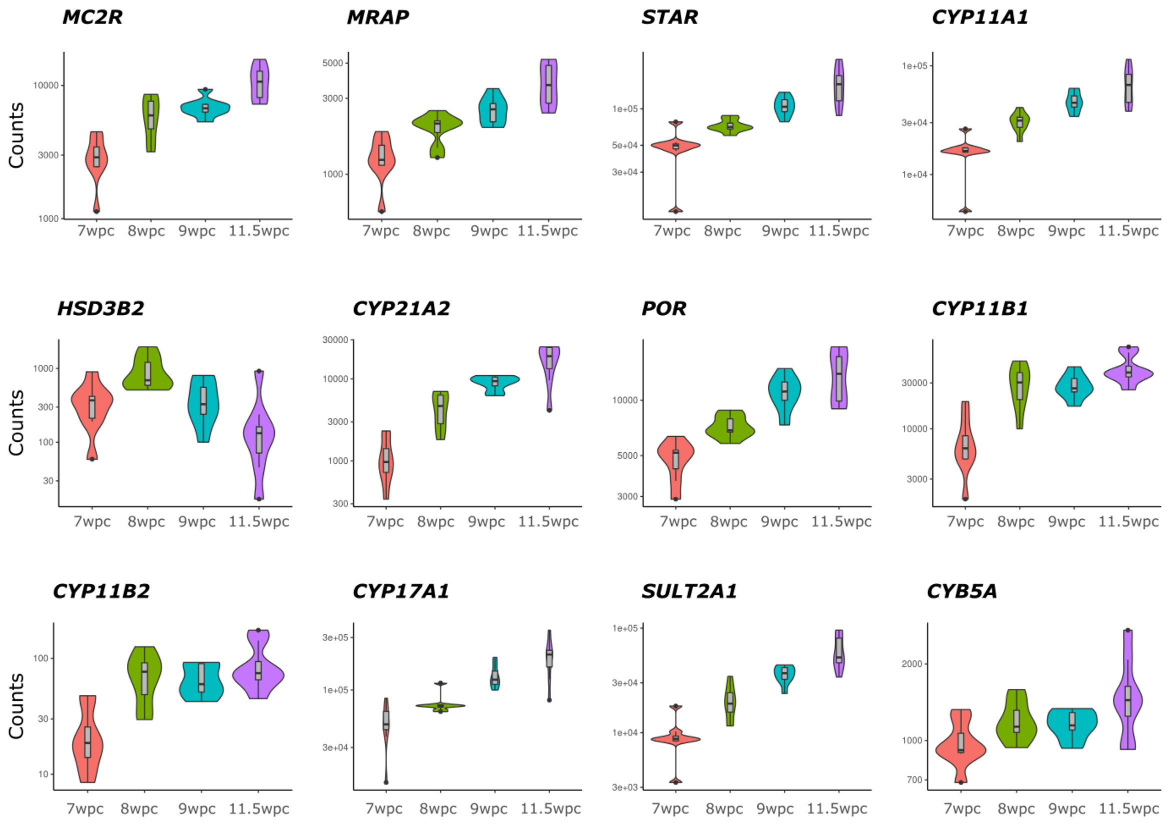


**Supplementary Fig. 6. Bulk RNA-seq time series changes in expression of key genes in the classic pathway of human adrenal steroidogenesis (see main Fig. 3d for reference).** Data (normalized counts) are shown as violin plots at each of the four different ages analyzed, with eight samples in each group.

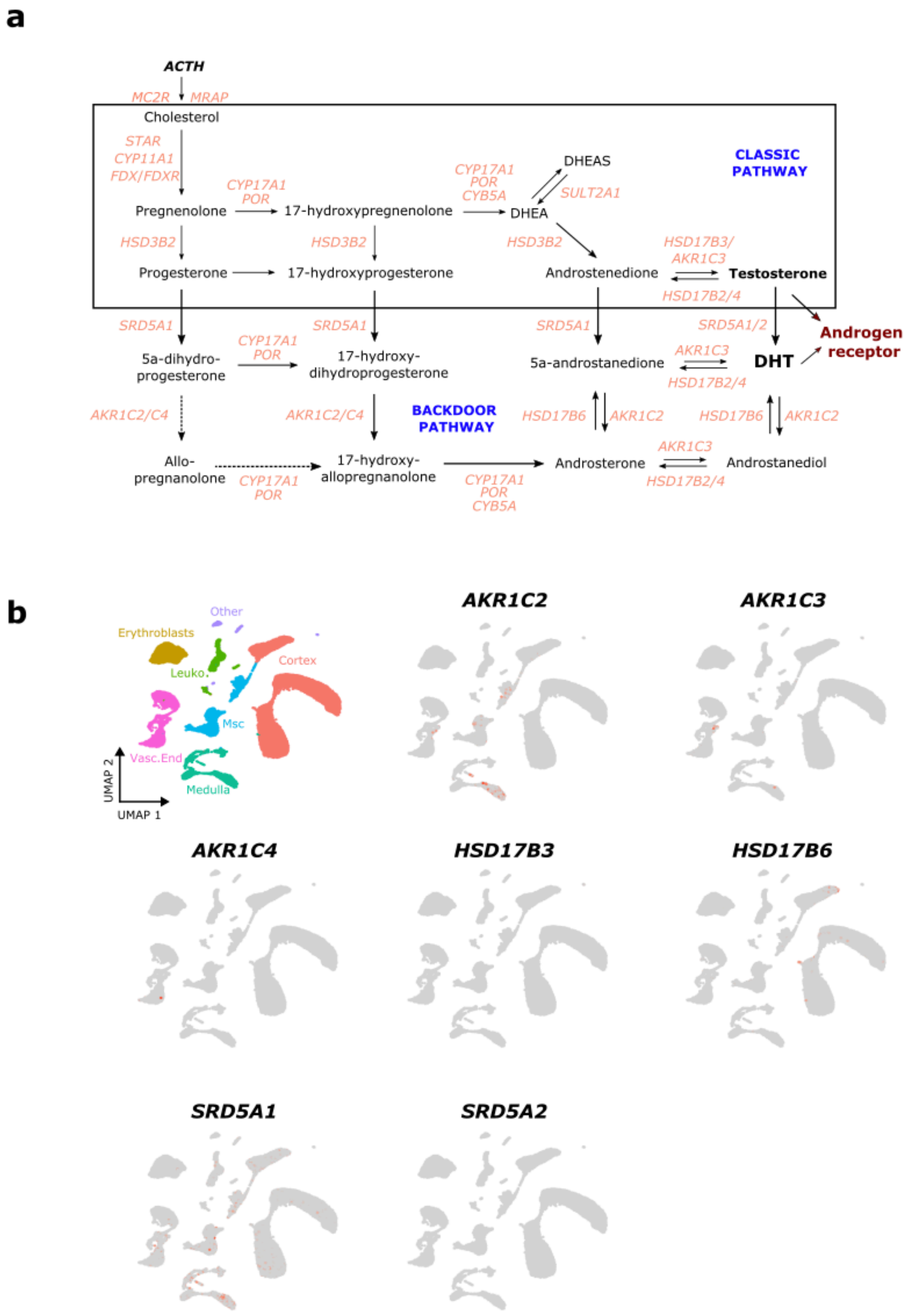


**Supplementary Fig. 7. Expression of "backdoor" pathway enzymes in the human fetal adrenal gland. a** Overview of the potential "backdoor" pathway to androgen (dihyhdrotestosterone, DHT) synthesis. **b** Feature plots showing single-cell RNA-seq expression (merged datasets) for key components of the "backdoor" pathway. Msc, mesenchyme.

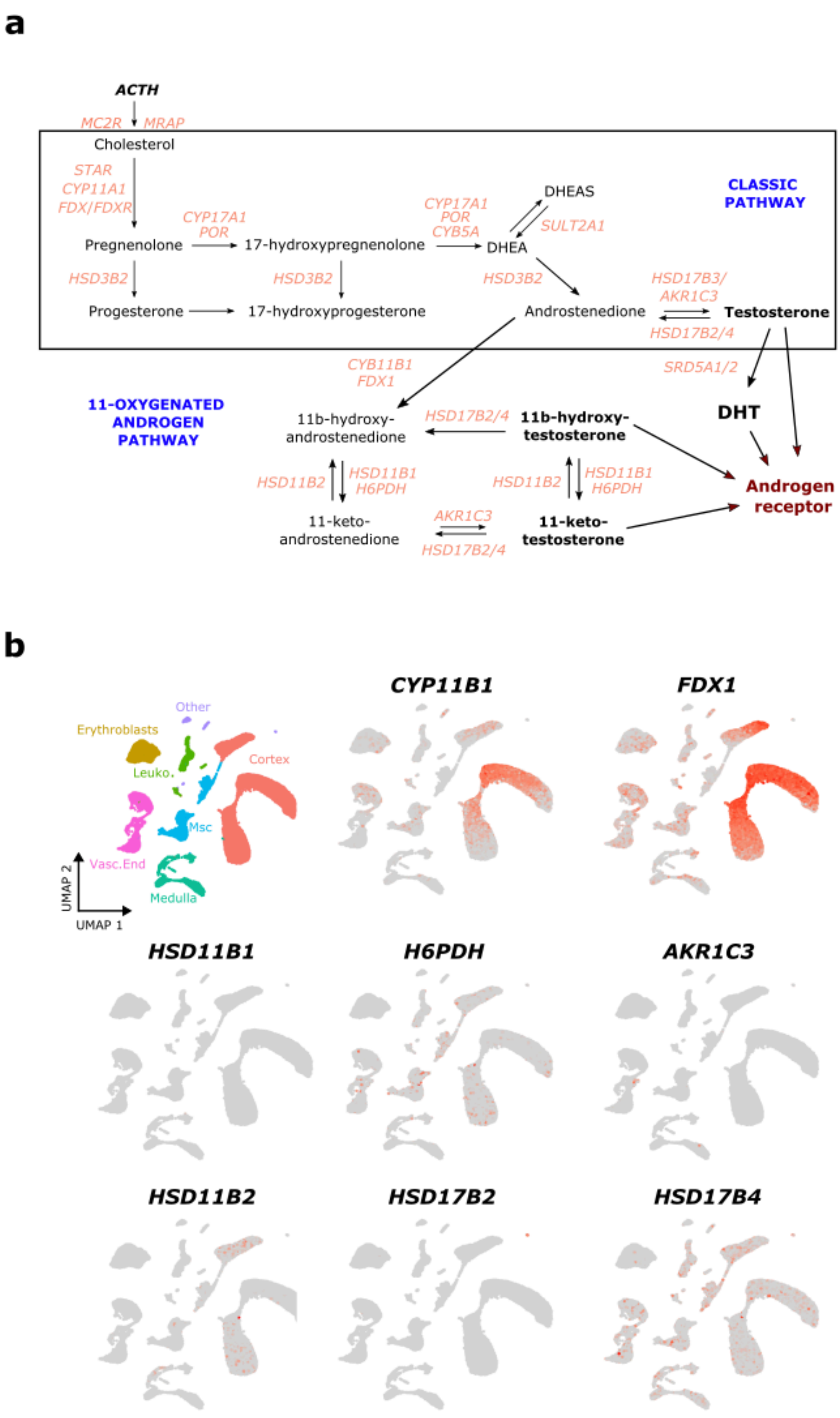


**Supplementary. Fig. 8. Expression of enzymes needed for 11-oxygenation of androgens in the human fetal adrenal gland. a** Overview of the potential “11-oxygenation” pathway. **b** Feature plots showing single-cell RNA-seq expression (merged dataset) for key components of the “11-oxygenation” pathway. Msc, mesenchyme.

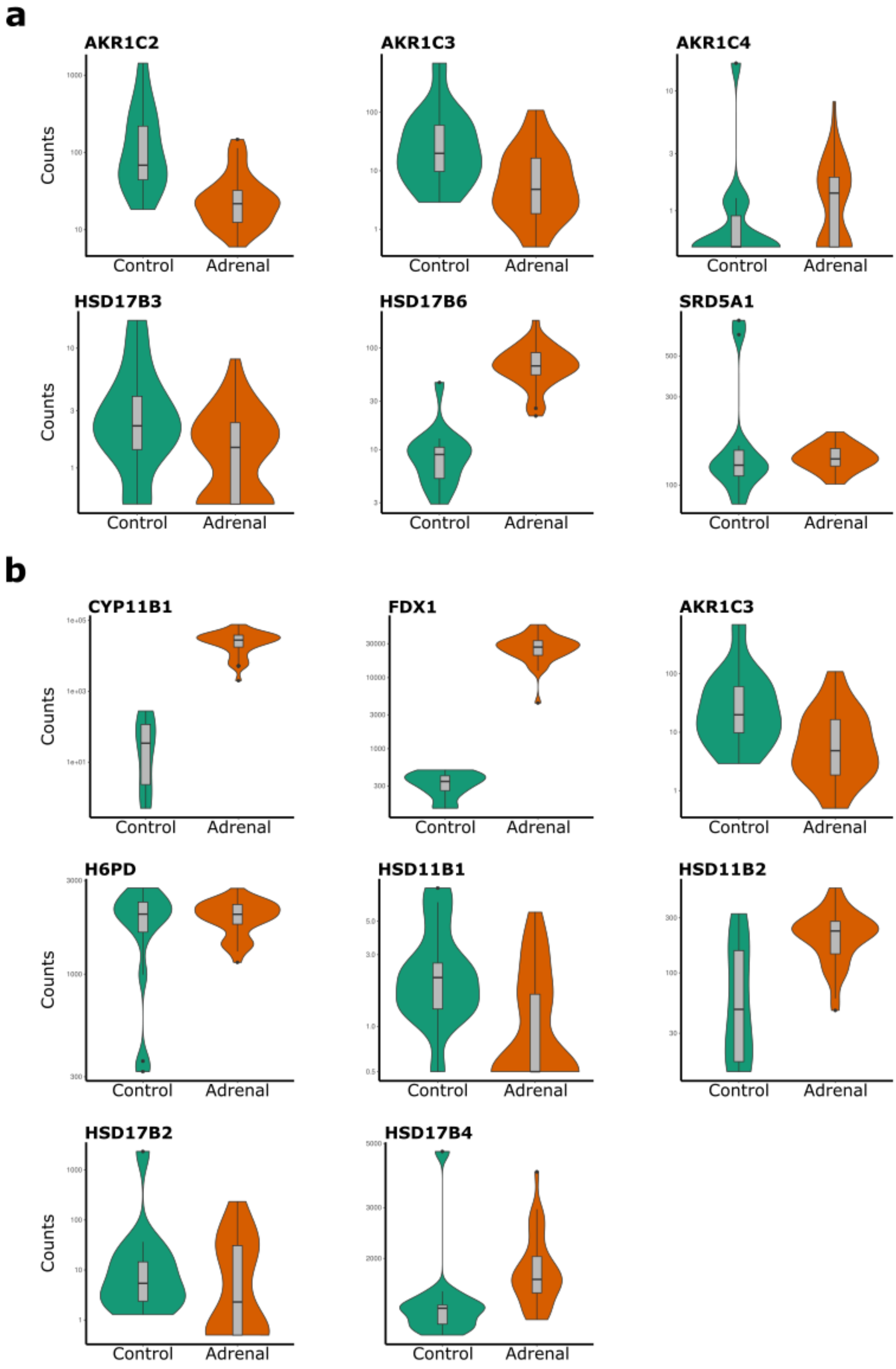


**Supplementary. Fig. 9. Expression of alternative pathway enzymes in bulk RNA-seq data for the adrenal glands compared to controls. a** Enzymes involved in the “backdoor” pathway. **b** Enzymes involved in 11-oxygenation of androgens. Data are shown as violin plots for normalized counts.

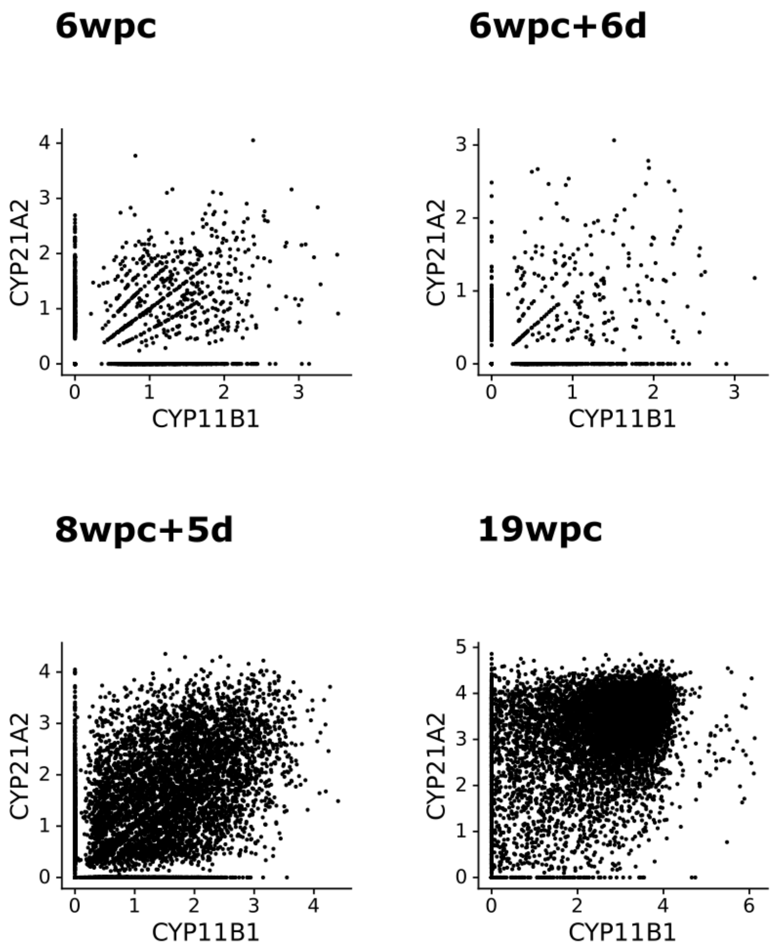


**Supplementary Fig. 10. Single-cell scatter plots showing expression of *CYP11B2* and *CYP21A2*.** Data from all four ages are shown, demonstrating an increase in expression of both genes with time.

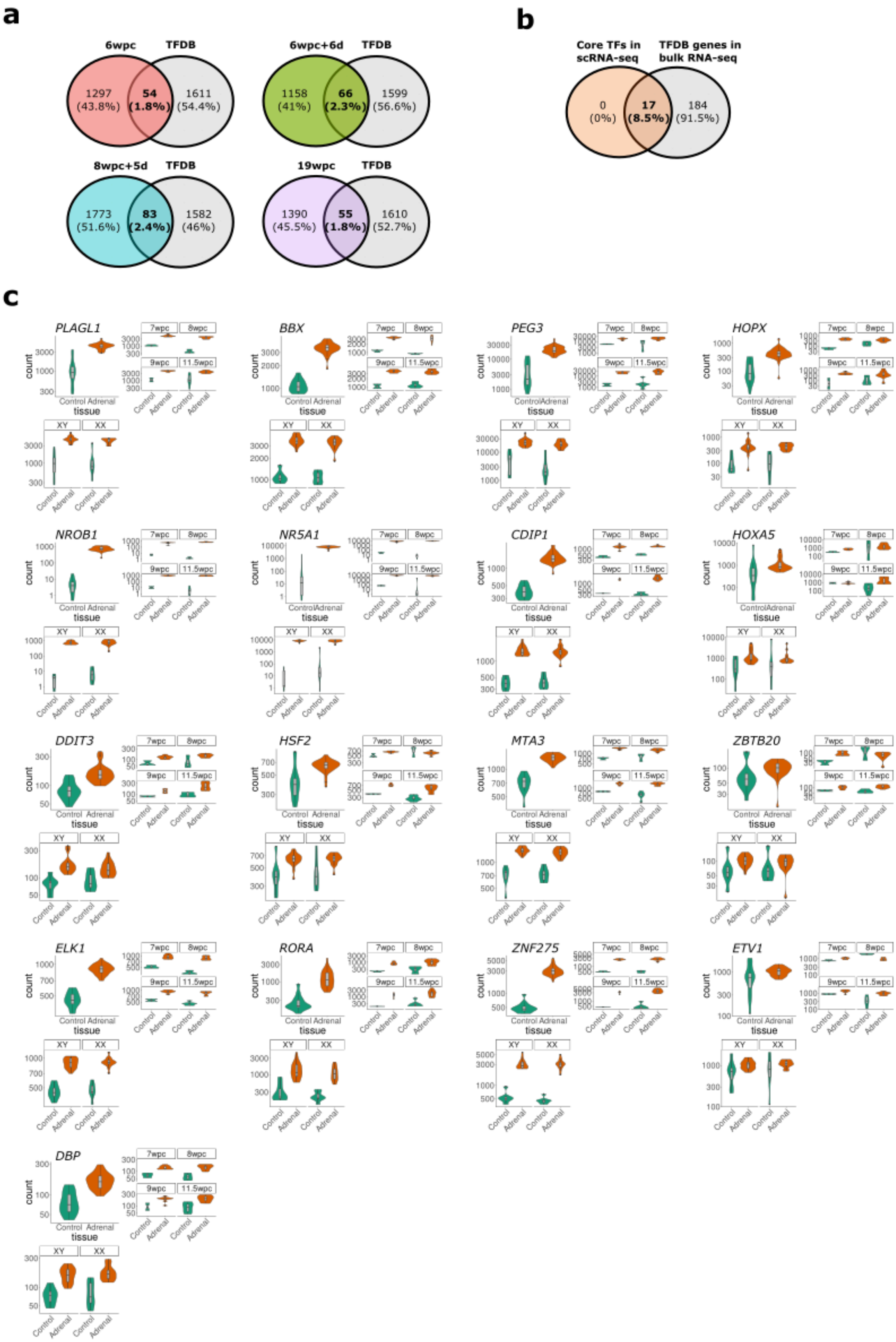
a
6wpc
TFDB
1297
(43.8%)
54
(1.8%)
1611
(54.4%)
6wpc+6d
TFDB
1158
(41%)
66
(2.3%)
1599
(56.6%)
8wpc+5d
TFDB
1773
(51.6%)
83
(2.4%)
1582
(46%)
19wpc
TFDB
1390
(45.5%)
55
(1.8%)
1610
(52.7%)
b
Core TFs in scRNA-seq
TFDB genes in bulk RNA-seq
0
(0%)
17
(8.5%)
184
(91.5%)
c
PLAGL1
BBX
PEG3
HOPX
NROB1
NR5A1
CDIP1
HOXA5
DDIT3
HSF2
MTA3
ZBTB20
ELK1
RORA
ZNF275
ETV1
DBP
count
tissue
Control
Adrenal
7wpc
8wpc
9wpc
11.5wpc
XY
XX

**Supplementary Fig. 11. Differentially-expressed transcription factors in the adrenal gland. a** Venn diagrams of transcription factors (transcription factor database, TFDB) differentially-expressed in the single-cell RNA-seq data (cortex cluster) at each age (log2 fold-change >0.25, padj<0.05). **b** Venn diagram showing the distribution of these 17 core transcription factors (TFs) identified within the scRNA-seq dataset (“Core TFs in scRNA-seq”) compared to TFDB genes that are differentially-expressed in the adrenal gland compared to controls using bulk-RNA seq (“TFDB bulk-RNA seq”) (adrenal>control, log10 fold change>2, padj<0.05). **c** Bulk RNA-seq normalized counts for core TFs showing expression compared to controls overall (*upper left individual panels*), expression compared to controls at each age (*upper right individual panels*) and expression in 46,XY and 46,XX samples (*lower left individual panels*).

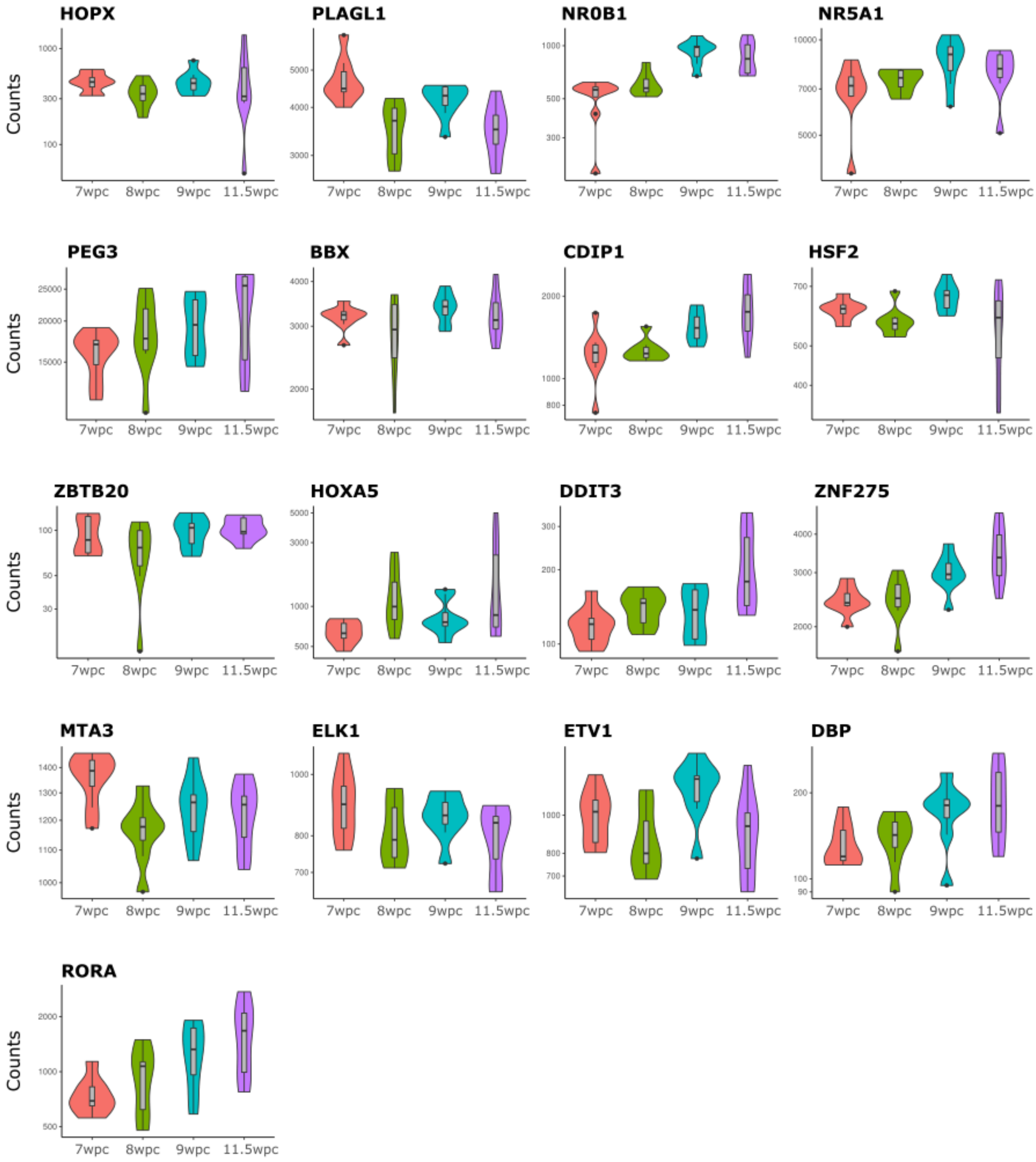


**Supplementary. Fig. 12. Bulk RNA-seq of "core" transcription factor expression at each age.** Data are shown as violin plots of normalized counts.

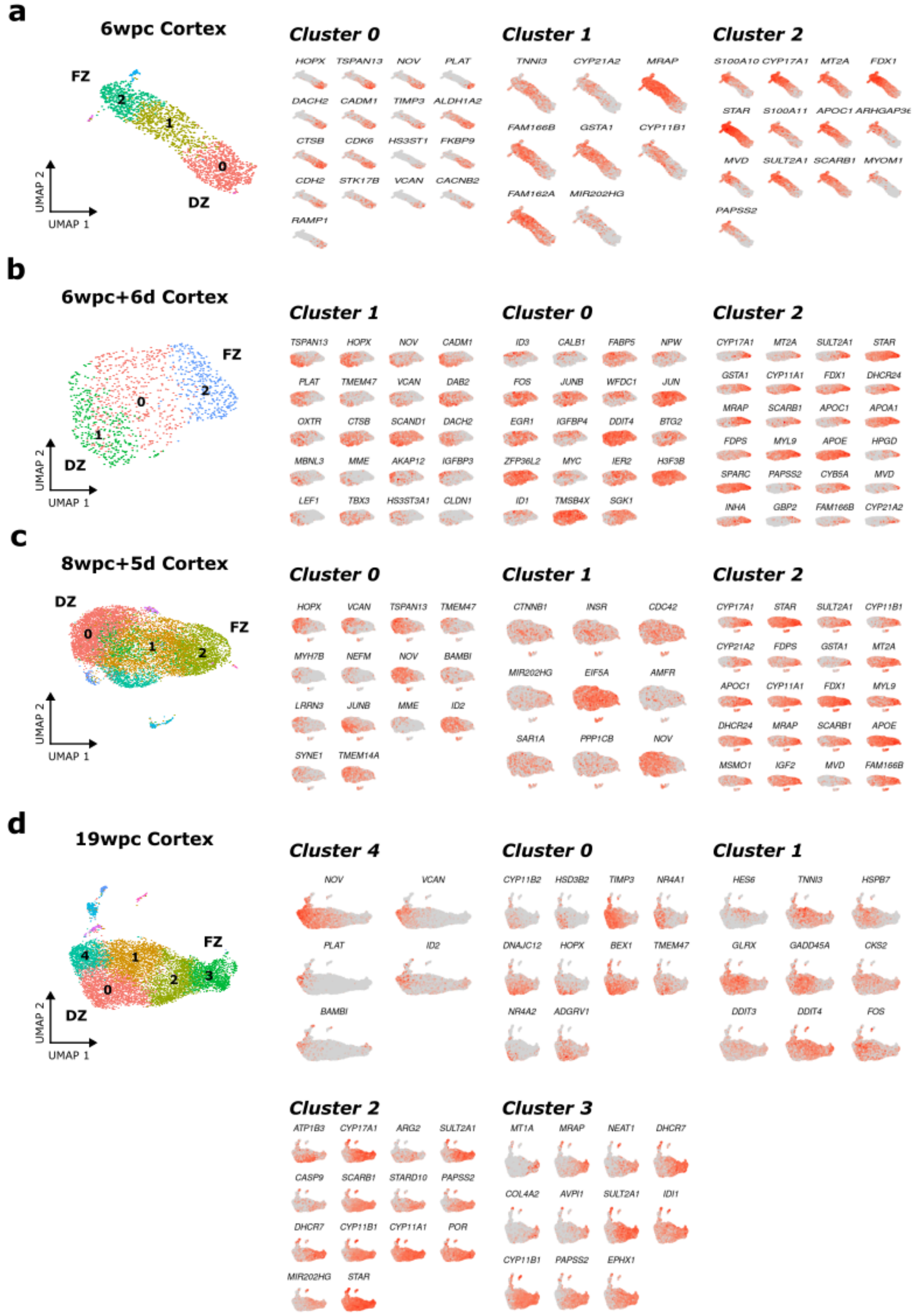


**Supplementary Fig. 13. UMAP and feature plots for scRNA-seq data showing the key differentially-expressed genes in each cluster identified at each age. a** 6wpc. **b** 6wpc+6d. **c** 8wpc+5d. **d** 19wpc. DZ, definitive zone; FZ, fetal zone. (See Supplementary Data 3 for complete list of gene expression data at each age)

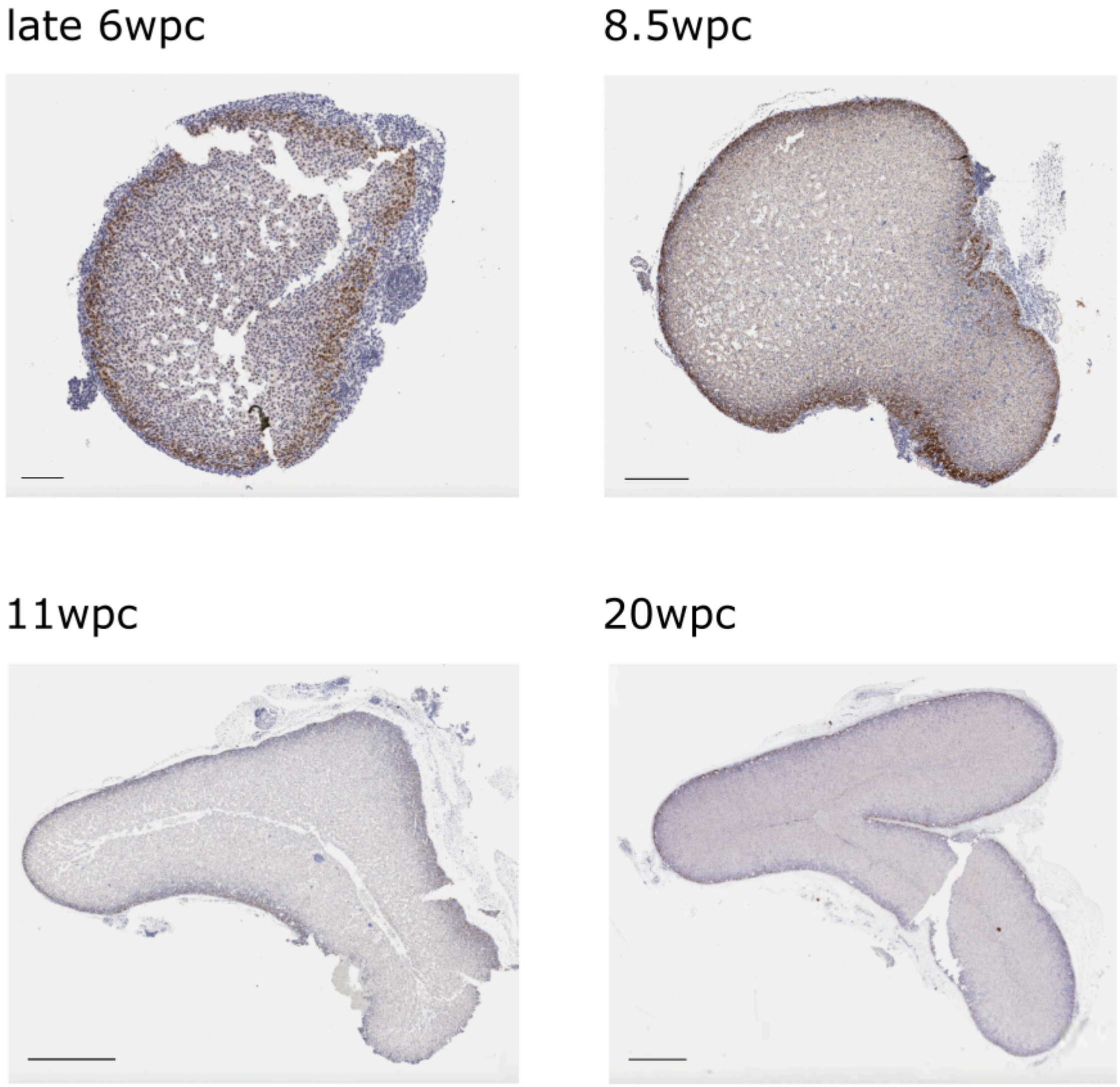


**Supplementary Fig. 14. Immunohistochemistry of HOPX in the human adrenal gland at different ages.** Scale bars: late 6wpc, 100μm; 8.5wpc, 400μm; 11wpc, 800μm; 20wpc, 1000μm.

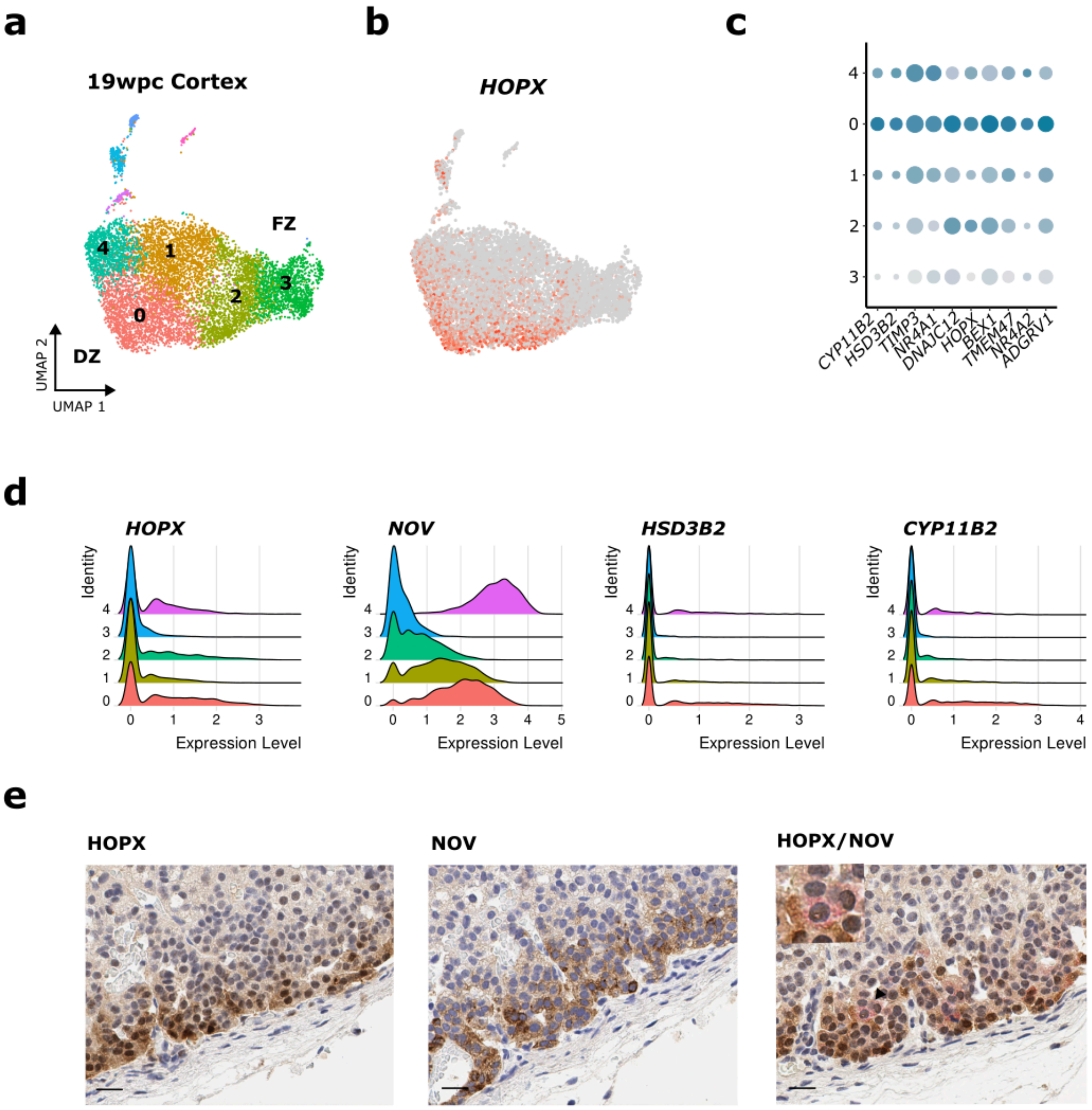


**Supplementary Fig. 15. Co-expression of *HOPX*/HOPX with related genes of interest in the 19wpc adrenal gland. a** UMAP of key clusters in scRNA-seq data. **b** Feature plot of *HOPX* expression. **c** Dot plot showing relative expression of the top 10 differentially-expressed genes in cluster 0 compared to other clusters (1, 2, 3, 4). **d** Ridge plots of *HOPX* expression and related genes of interest in cluster 0 and cluster 4. **e** Immunohistochemistry of HOPX expression (brown peripheral nuclear staining, *left panel*), NOV expression (brown cytoplasmic staining into deeper layers, *center panel*), and dual staining (HOPX, brown; NOV, pink, *right panel*). Scales all 20µm.

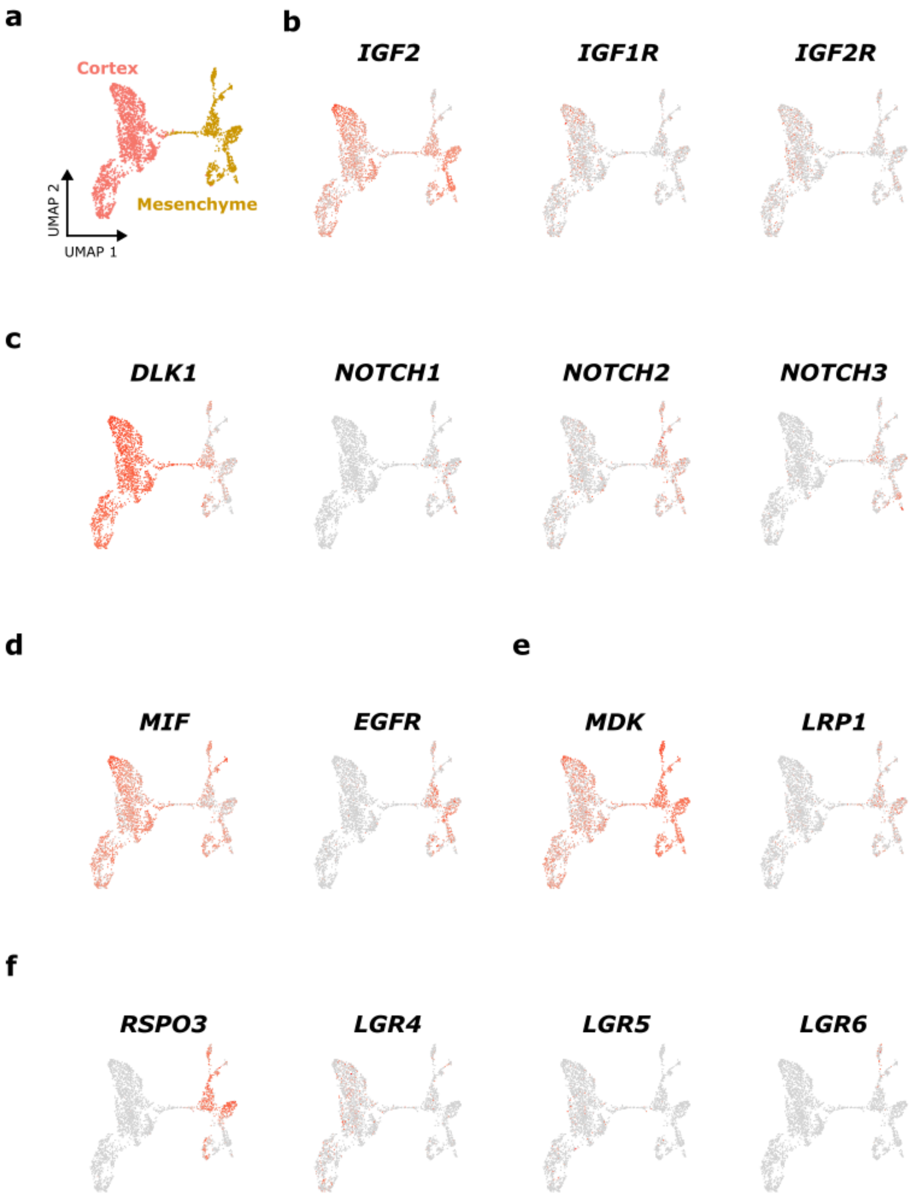


**Supplementary Fig. 16. Expression of key genes involved in ligand-receptor interactions in the mesenchyme and adrenal cortex clusters at 6wpc+6d identified by cell-cell communication network analysis.** Data relates to main Fig. 6b. **a** Reference UMAP showing mesenchyme and adrenal cortex clusters. **b** Feature plot showing expression of *IGF2* (ligand) and related cognate receptors. **c** *DLK1* (ligand) and *Notch* family receptors. **d** *MIF* (ligand) and *EGFR*. **e** *MDK* (ligand) and *LRP1*. **f** *RSPO3* (ligand) and related *LGR* receptors.

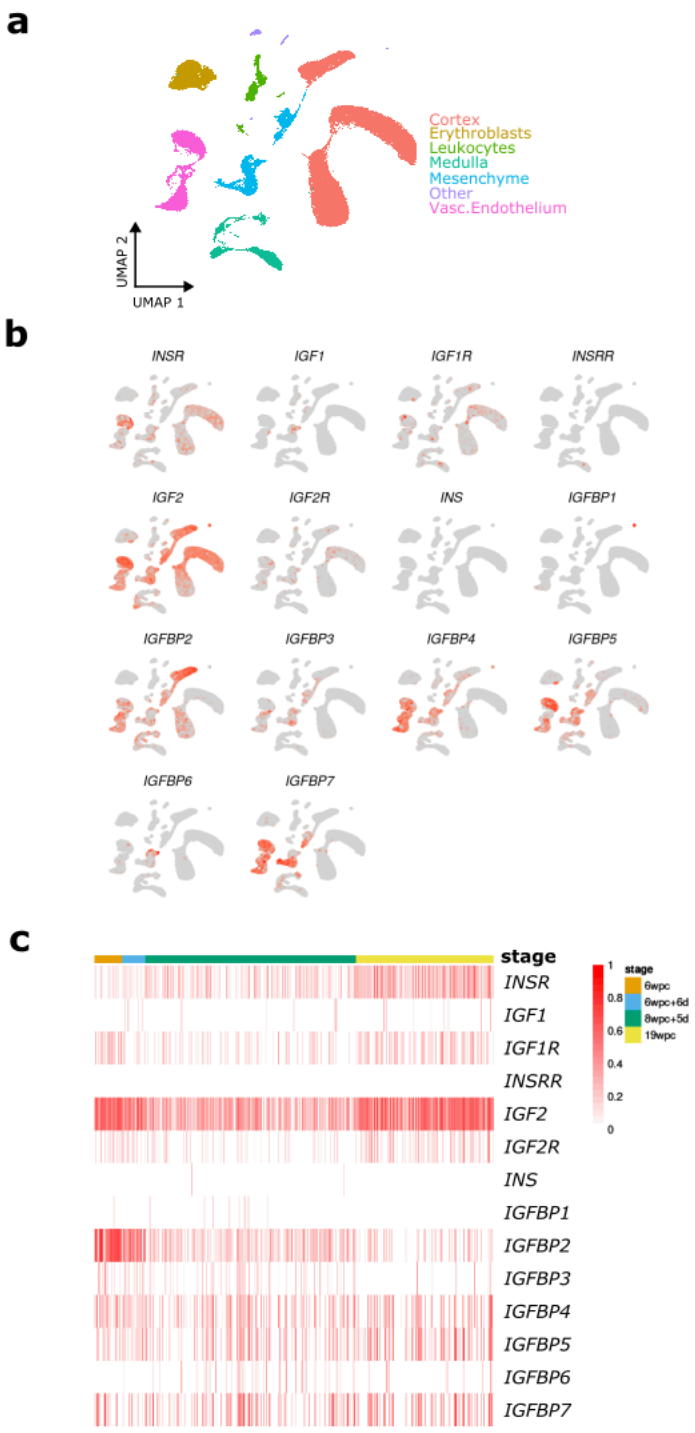


**Supplementary Fig. 17. scRNA-seq expression of insulin-like growth factor II (*IGF2*) and related genes in the adrenal gland. a** Reference merged UMAP (as shown in Fig. 1g). **b** Feature plot for all clusters in the adrenal gland showing expression of *IGF2* and related genes. **c** Heatmap of expression of *IGF2* and related genes in the adrenal cortex cluster across the four ages studied.

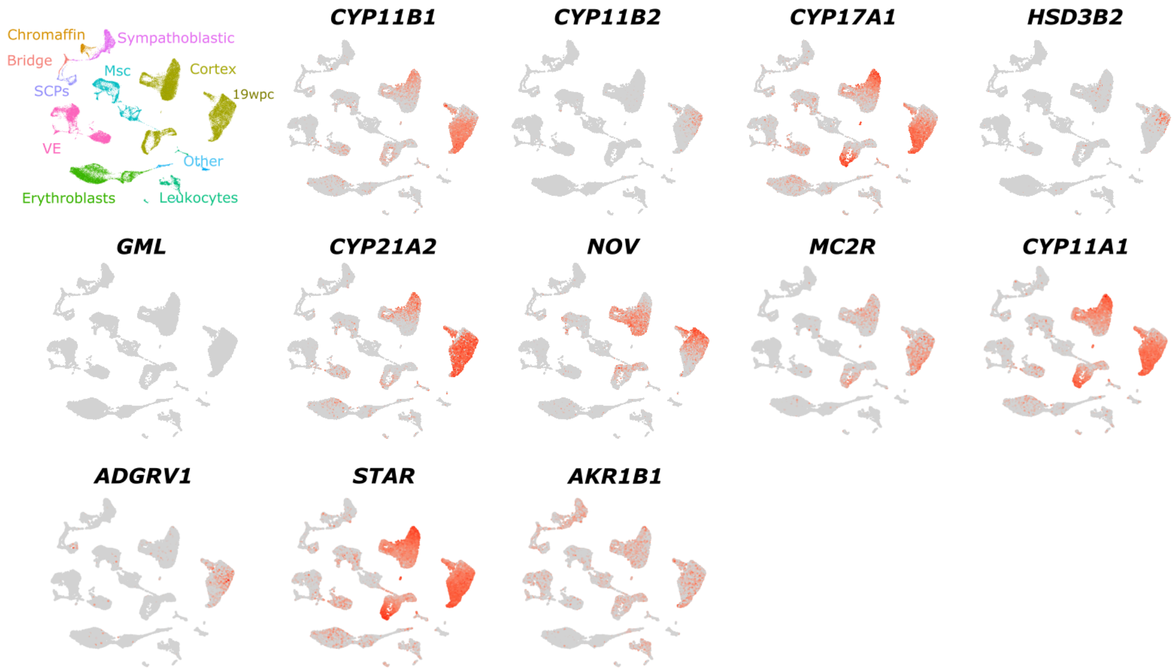


**Supplementary Fig. 18. Feature plots showing fetal adrenal expression of genes corresponding to the genes (n=12) that are most highly differentially expressed in the adult adrenal gland (Human Protein Atlas, www.proteinatlas.org).** Data refer to the dot plot in Main Fig. 8a, but show the increase in expression of several genes with age (19wpc cluster indicated)
